# The developmentally-timed decay of an essential microRNA family is seed sequence-dependent

**DOI:** 10.1101/2021.11.19.469346

**Authors:** Bridget F. Donnelly, Bing Yang, Acadia L. Grimme, Karl-Frédéric Vieux, Chen-Yu Liu, Lecong Zhou, Katherine McJunkin

## Abstract

MicroRNA (miRNA) abundance is tightly controlled by regulation of biogenesis and decay. Here we show that the *mir-35* miRNA family undergoes selective decay at the transition from embryonic to larval development in *C. elegans*. The seed sequence of the miRNA is necessary and largely sufficient for this regulation. Sequences outside the seed (3’ end) regulate *mir-35* abundance in the embryo but are not necessary for sharp decay at the transition to larval development. Enzymatic modifications of the miRNA 3’ end are neither prevalent nor correlated with changes in decay, suggesting that miRNA 3’ end display is not a core feature of this mechanism and further supporting a seed-driven decay model. Our findings demonstrate that seed sequence-specific decay can selectively and coherently regulate all redundant members of a miRNA seed family, a class of mechanism that has great biological and therapeutic potential for dynamic regulation of a miRNA family’s target repertoire.

## Introduction

microRNAs (miRNAs) are small non-coding RNAs (∼22-23 nucleotides) that, when bound by Argonaute, form the miRNA-induced silencing complex (miRISC) and negatively regulate target mRNAs (Dallaire et al., 2018). The biogenesis of miRNAs has been well described; first, miRNAs are transcribed as a primary miRNA (pri-miRNA): a long transcript containing a ∼35 base pair stem-loop structure formed by intramolecular base-pairing (Fang and Bartel, 2015; Han et al., 2006; Ma et al., 2013; Zeng et al., 2005). The double-stranded RNA hairpin structure of the pri-miRNA is recognized by the Microprocessor complex (Drosha and DGCR8/Pasha) (Fang and Bartel, 2015; Han et al., 2006; Ma et al., 2013; Zeng et al., 2005). The catalytic RNase III domain of Drosha cleaves the pri-miRNA, resulting in the generation of a miRNA precursor (pre-miRNA) hairpin structure with a two-nucleotide overhang at the 3’ end (Denli et al., 2004; Gregory et al., 2004; Han et al., 2004; Landthaler et al., 2004). Once exported from the nucleus, the pre-miRNA is cleaved by the RNase III enzyme Dicer into a ∼22-23 nucleotide duplex that is loaded into Argonaute (Bernstein et al., 2001; Grishok et al., 2001; Hutvágner et al., 2001; Ketting et al., 2001; Knight and Bass, 2001). The mature guide strand remains in the Argonaute protein, becoming a part of the miRISC, while the star strand is ejected and degraded (Iwasaki et al., 2010, 2015). Once the mature miRNA is incorporated into miRISC, miRISC uses the bound miRNA guide strand to target complementary regions in the 3’UTR of mRNAs to silence gene expression by translational repression, deadenylation and decay of the target mRNA (Dexheimer and Cochella, 2020).

The interaction between the miRNA and mRNA target is primarily mediated through nucleotides 2-8 at the 5’ end of the miRNA (Brennecke et al., 2005; Lewis et al., 2003). This region, called the seed sequence, is the defining characteristic of a miRNA family, a group of miRNAs that act largely redundantly on an overlapping set of target genes due to their identical seed sequences (Alvarez-Saavedra and Horvitz, 2010; Parchem et al., 2015). Additional supplemental base pairing between the 3’ end of the miRNA and the target RNA occurs in some cases, conferring some differences in target repertoire of miRNA family members (which share a seed sequence but may differ in their 3’ end sequences) (Brancati and Großhans, 2018; Broughton et al., 2016; Helwak et al., 2013; Ye Duan, Isana Veksler-Lublinsky, 2021).

While much is known about the biogenesis and functions of miRNAs, relatively little is known about the mechanisms of decay of mature miRNAs. While half-lives of miRNAs vary, what determines these differences in stability is for the most part unknown (Bail et al., 2010; Kingston and Bartel, 2019; Lehrbach et al., 2012; Marzi et al., 2016; Miki et al., 2014; Reichholf et al., 2019; Vieux et al., 2021). Thus far, multiple phenomena regulating miRNA stability have been observed, with different degrees of sequence-specificity.

Some decay pathways appear to be largely independent of miRNA sequence. In *C. elegans*, the 5’ to 3’ nuclease XRN-2, along with DCS-1, maintain wild type miRNA levels by degrading many (though not all) miRNAs (Bossé et al., 2013; Chatterjee et al., 2009). In the mouse retina, an undefined mechanism induces decay of most miRNAs upon light-dependent neuronal activity (Krol et al., 2010). At the maternal to zygotic transition in *Drosophila*, terminal adenylation of maternal miRNAs by the noncanonical poly(A) polymerase, Wispy, induces their wholesale clearance (Lee et al., 2014). In other species, 3’ nucleotide addition (tailing) has also been proposed to destabilize miRNAs in a sequence-independent manner (Boele et al., 2014; Katoh et al., 2015; Knouf et al., 2013; Lee et al., 2019; Shukla et al., 2019; Wyman et al., 2011; Yang et al., 2020a).

Other miRNA decay pathways are guided by moderate sequence-specificity. One example is Tudor SN-mediated miRNA decay (TumiD) (Elbarbary et al., 2017a, 2017b). In TumiD, the endonuclease Tudor-SN (TSN) cleaves a few dozen miRNAs at CA and UA dinucleotides greater than 5 nucleotides away from the 3’ and 5’ ends of the miRNA (Elbarbary et al., 2017a, 2017b). A more specific phenomenon confers instability to several members of the extended miR-16 family; this decay is dependent on sequences in the both the seed and the 3’ portion of the miRNA (Rissland et al., 2011). Rapid decay of miR-382 is dependent on the 7 nucleotides at the 3’ end of the miRNA (Bail et al., 2010).

The most sequence-specific mechanism of miRNA decay is called target-directed miRNA degradation (TDMD). TDMD occurs when a high-abundance RNA (the TDMD “trigger”) binds to a miRNA with extensive complementarity to both the seed sequence and the 3’ half of the miRNA (Ameres et al., 2010; Baccarini et al., 2011; Bitetti et al., 2018; Cazalla et al., 2010; Ghini et al., 2018; Kleaveland et al., 2018; Libri et al., 2012; Marcinowski et al., 2012; la Mata et al., 2015; Piwecka et al., 2017). This extensive base pairing induces a conformational change that pulls the 3’ end of the miRNA out of the PAZ domain of Argonaute, making it accessible to modification by untemplated nucleotide additions (tailing) and exonucleolytic cleavage (trimming) (Sheu-Gruttadauria et al., 2019; Yang et al., 2020a). Recently, the Cullin-RING E3 ubiquitin ligase ZSWIM8 was identified as an effector of TDMD, leading to the model that the Argonaute and/or RNA conformation induced by extensive base pairing is recognized by ZSWIM8 for ubiquitylation and subsequent decay of the miRNA:Argonaute complex (Han et al., 2020; Shi et al., 2020).

The regulation of miRNA expression during development is crucial to ensure properly timed developmental transitions, but the extent to which miRNA decay contributes to ensuring proper temporal expression patterns of miRNAs and how development is coupled to timing of decay are not known.

In this work, we set out the examine the mechanism of decay of the *mir-35* family of miRNAs. The *mir-35* family consists of 8 miRNAs, *mir-35-42* (Alvarez-Saavedra and Horvitz, 2010). The *mir-35* family members are maternally contributed as well as zygotically expressed in early embryogenesis, and they are sharply degraded at the transition from embryo to the first larval stage, L1 (Stoeckius et al., 2009; Wu et al., 2010). Understanding the mechanism of this decay will shed light on how selective miRNA decay occurs and how it is coupled to development.

The *mir-35* family is one of two miRNA seed families that are necessary for *C. elegans* embryogenesis. Because of their identical seed sequences, the *mir-35* family members are functionally redundant; deletion of any single miRNA has no detectable phenotypic consequences, whereas deletion of the whole family results in embryonic lethality (Alvarez-Saavedra and Horvitz, 2010). The *mir-35* family miRNAs also play multiple roles in development, including promoting maximal fecundity, ensuring sex determination, and regulating cell death (Doll et al., 2019; Flamand et al., 2016, 2017; Kagias and Pocock, 2015; Liu et al., 2011; Massirer et al., 2012; McJunkin and Ambros, 2014, 2017; Tran et al., 2019; Yang et al., 2020b; Zhao et al., 2019).

How the *mir-35* family is targeted for selective decay at the end of embryogenesis is not known. A recent study showed that the TDMD factor ZSWIM8 (known as EBAX-1 in *C. elegans*) drives instability of the *mir-35* family, suggesting that the *mir-35* family is subject to TDMD (Shi et al., 2020; Wang et al., 2013). However, positions of the miRNA that are usually involved in the base-pairing interactions that drive TDMD (in the 3’ half of the miRNA) are highly degenerate across the *mir-35* family members (Figure 1A), suggesting that the mechanism of *mir-35* decay may differ from previously-described examples of TDMD, and may represent a novel type of selective miRNA decay mechanism.

**Figure 1.**
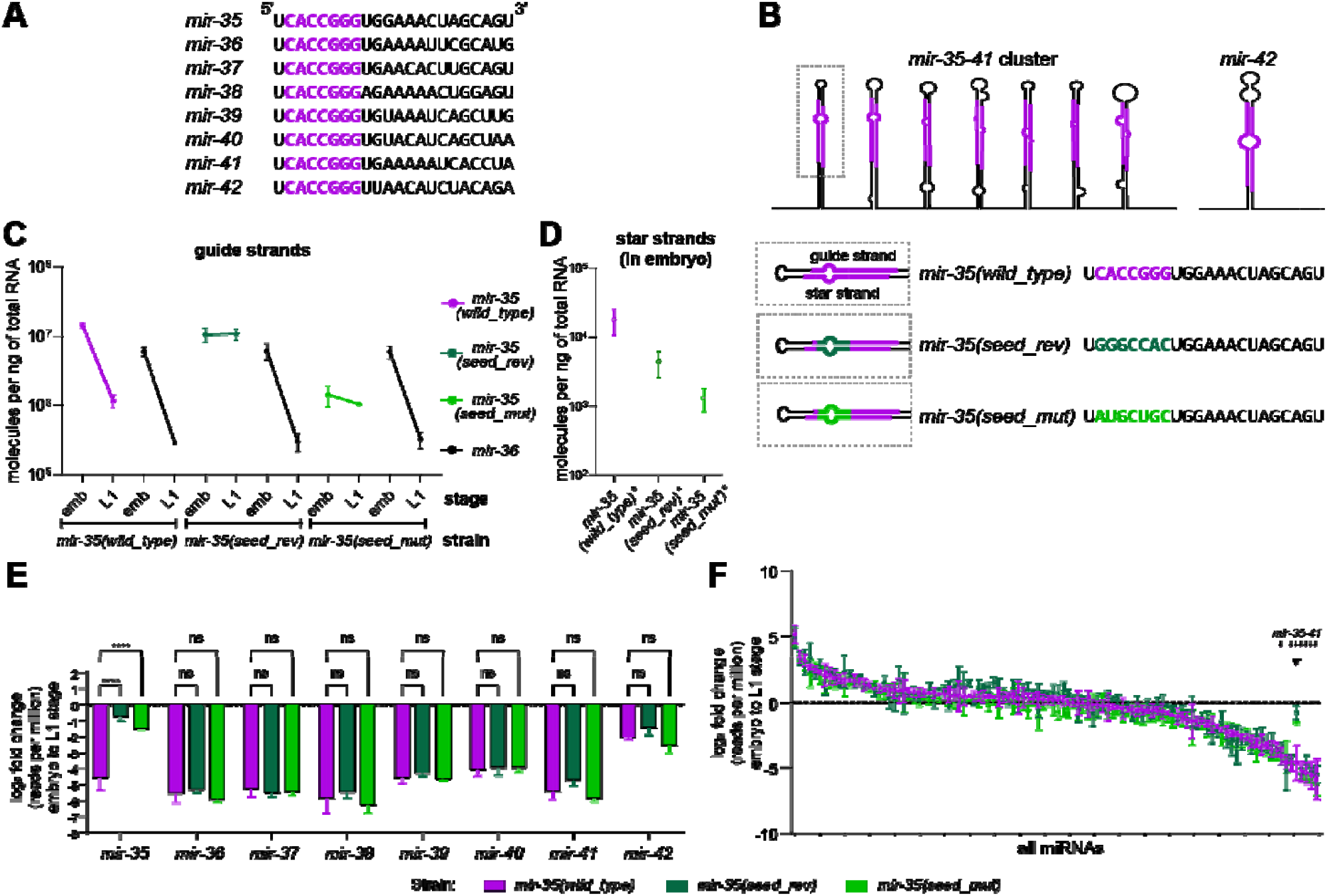
*mir-35* decay is seed sequence-dependent. (A) Sequences of *mirs-35-42*. The identical seed sequence is shown in purple. (B) Schematic of the *mir-35-41* cluster and *mir-42* showing the structure of the hairpins in the primary transcripts. The guide and star strands are highlighted in purple (top). Below is the wild type *mir-35* sequence (purple) and the two seed mutant *mir-35* variants (green). Both seed mutant *mir-35* variants have compensatory mutations to the star strand to maintain the wild type-like structure of the mutant *mir-35* hairpin. (C) Absolute quantification of *mir-35* and *mir-36* in embryos and L1. For some error bars, the range is smaller than the symbol. (D) Absolute quantification of star strands of *mir-35* and mutant variants in embryos. (E-F) Log_2_(fold change) in abundance from embryo to L1, calculated from normalized deep sequencing reads for either the *mir-35-42* family (E) or all miRNAs with >50 RPM in wild type (F). Note that color of bar indicates strain, not necessarily a mutant miRNA; only *mir-35* is mutated in the indicated mutant strains. (E) Two-way ANOVA was performed, followed by Dunnett’s multiple comparisons test. ****p-value < 0.0001 Small arrows in (F) indicate positions of *mir-35-41* on ranked x-axis, and arrowhead indicates *mir-35* with mutant variants. (C-F) Mean and SEM of three biological replicates are shown.

Here we show that the *mir-35* family is regulated at the level of decay at the embryo to L1 transition in *C. elegans*. We demonstrate that the seed sequence of *mir-35* is necessary and largely sufficient for this developmentally timed decay. This decay is not correlated with high levels of miRNA 3’ tailing and trimming. Together, these data suggest that this miRNA family is regulated by a distinct – but possibly related – mechanism to TDMD. Seed-specific decay mechanisms such as this are likely to be more widespread in biological systems since they have potential to co-regulate all members of a redundant miRNA family, potentially allowing dynamic derepression of the miRNA family’s target genes.

## Results

### *mir-35* decay is seed sequence-dependent

The *mir-35* family is selectively decayed at the embryo to L1 transition (Stoeckius et al., 2009; Wu et al., 2010). We wondered if its decay is a selectively regulated process or, alternatively, just a result of transcriptional shutoff in late embryogenesis. We also wondered whether its characteristic seed sequence played a role in this putative selectively regulated decay. To this end, we used CRISPR/Cas9 to mutate the genomic locus which encodes seven of the eight *mir-35* family members, *mir-35-41*, on a single transcript (Figure 1B). (*mir-42* is clustered with unrelated miRNAs at a different genomic locus.) We made targeted mutations to the seed sequence of the first hairpin in the *mir-35-41* cluster (*mir*-*35*) using CRISPR/Cas9. This approach leaves the remainder of the *mir*-*35-41* cluster intact, which serves two purposes: 1) *mir-35* loss-of-function phenotypes are not induced since the other family members remain wild type, and 2) *mir-36-41* serve as internal controls derived from the same transcript as *mir-35*. Both strands of the hairpin encoding *mir-35* were mutated to preserve secondary structure and support efficient processing (Figure 1B). One of the mutations was a reversal of the seed sequence, referred to hereafter as *mir-35(seed_rev)*), whereas the other mutation replaced the *mir-35* seed sequence with random nucleotides (*mir-35(seed_mut)*) (Figure 1B). (See Methods and Tables S1 and S2 for all allele, strain, and genome editing details.)

To determine if biogenesis of *mir-35* was affected by these seed mutations, we quantified *mir-35* and the mutant variants using miRNA-Taqman qPCR, along with synthetic RNA oligonucleotides to generate standard curves for absolute quantification (Figure S1). The embryo concentration of *mir-35(seed_rev)* is similar to wild type *mir-35* (0.7-fold change), while *mir-35(seed_mut)* is ten-fold lower (Figure 1C). To determine if the changes in the level of the *mir-35* variants were at the level of biogenesis or post-biogenesis, we examined the abundance of their star strands in the embryo. Changes is abundance of the *mir-35* variant star strands is similar to those in the respective guide strands (Figure 1D); these coupled changes suggest that the decreased abundance of *mir-35(seed_mut)* is due to loss of efficiency in biogenesis.

Next, we examined whether the decay of *mir-35* at the embryo to L1 transition is altered by seed mutations. (Note that, because we use arrested L1 samples, post-embryogenesis growth has not begun, so any decreases in miRNA abundance must be attributed to decay rather than dilution caused by growth.) As expected, we observed a strong reduction in wild type *mir-35* at the transition from embryo to L1, with 12-fold lower abundance in L1 (Figure 1C). However, the decay of mutant *mir*-*35*(*seed_rev*) and *mir*-*35*(*seed_mut*) at the transition from embryo to L1 was greatly attenuated to essentially no change and 1.3-fold lower in L1 than embryo, respectively (Figure 1C). *mir-35(seed_rev)* derived from a second CRISPR allele with altered precursor secondary structure also showed attenuated decay (Figure S2). Therefore, the decay of *mir-*35 depends on its seed sequence. Importantly, the decay of *mir-36* was not altered by the mutations in *mir-35* (Figure 1C). This decoupling of the behavior of mutant *mir-35* and wild type *mir-36* – which share a primary transcript – further shows that *mir-35* family decay is regulated post-transcriptionally.

To confirm and extend these findings, we performed deep sequencing to profile all miRNAs. Deep sequencing confirmed that *mir-35* was the only miRNA altered in abundance by these mutations in embryo or L1 samples (Figure S3, Tables S3 and S4). Consistent with the qPCR, wild type *mir-35* displayed sharp decay at the embryo to L1 transition, and the *mir-35* seed mutants were resistant to this decay (Figure 1E, Table S4). The decay of the other members of the *mir-35* family was not affected by *mir-35* seed mutations, despite most members sharing the *mir-35-41* primary transcript (Figure 1E). Global analysis further demonstrated the selectivity of the decay of the *mir-35* family in this developmental window: *mir-35-41* represent seven of the eight most sharply downregulated miRNAs at this time point in wild type (Figure 1F, Table S4). This analysis also reiterates the specificity of the effect of the *mir-35* seed mutations, further demonstrating that the regulation of other miRNAs is not affected by these mutations (Figure 1F).

Together these results show that the decay of the *mir-35* family at this developmental transition is a selectively regulated decay process (rather than simply the result of synchronous decay after transcriptional shutoff), since the behavior of miRNAs derived from the same transcript can be de-coupled. Furthermore, these results show that the *mir-35* seed sequence is required for this regulated decay.

### *mir-35* 3’ end mutants undergo efficient decay at the embryo to L1 transition

The necessity of the seed sequence for *mir-35* decay (Figure 1) and the recent implication of the TDMD factor ZSWIM8/EBAX-1 in regulating stability of the *mir-35* family (Shi et al., 2020) together suggest a TDMD-like decay mechanism. However, the degeneracy of sequences in the 3’ region of the miRNA across the *mir-35* family members (Figure 1A) suggests that the mechanism may differ from previous examples of TDMD since multiple trigger RNAs would be necessary to bind with extensive complementarity to all the family members. (Note that results from *mir-35(seed_rev)* and *mir-35(seed_mut)* rule out an antisense RNA from the *mir-35-41* cluster acting as a TDMD trigger RNA, since mutations at the genomic locus would not disrupt base-pairing with an antisense transcript.)

Therefore, we next investigated whether the 3’ portion of the miRNA plays a role in *mir-35* family decay and if the *mir-35* seed sequence is sufficient for decay. To test this, we used CRISPR to generate two *mir-35* mutant strains in which the non-seed (hereafter referred to as the 3’ end) residues of the miRNA are mutated. The first mutant is comprised of the *mir-35* seed sequence with a 3’ end containing nucleotides that are either not present or rare among all *mir-35* family members at a given position, while preserving overall GC content (*mir-35(mut_3’)*) (Figure 2A). The second mutant is a *mir-35/mir-82* hybrid composed of the *mir-35* seed sequence and *mir-82* 3’ end (*mir-35(mir-82_3’)*) (Figure 2A). The *mir-82* sequence was chosen for the non-seed region of *mir-35* because *mir-82* expression is steady rather than downregulated at the embryo to L1 transition (Kato et al., 2009).

**Figure 2.**
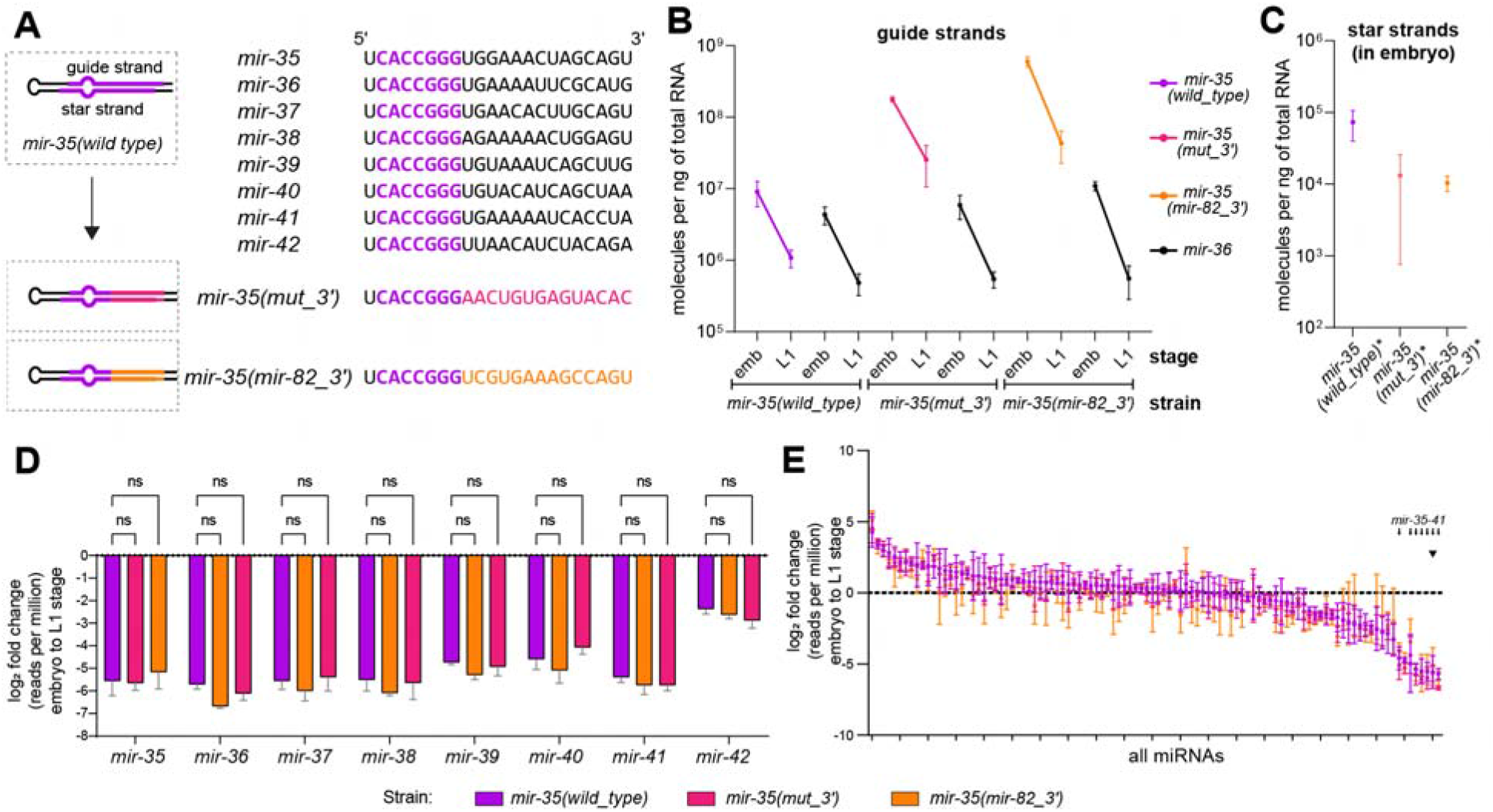
*mir-35* 3’ end mutants do not alter decay. (A) Sequences of *mirs-35-42* with the identical seed sequences shown in purple (top). Schematic of the *mir-35* 3’ end mutants (bottom). Both 3’ end mutants have compensatory mutations to the star strand to maintain the wild type like structure of the mutant *mir-35* hairpin. (B) Absolute quantification of *mir-35* and *mir-36* in embryos and L1. Mean and SEM of two to three biological replicates are shown. (C) Absolute quantification of star strands of *mir-35* and mutant variants in embryos. (D-E) Log_2_(fold change) in abundance from embryo to L1, calculated from normalized deep sequencing reads for either the *mir-35-42* family (D) or all miRNAs with >50 RPM in wild type (E). Note that color of bar indicates strain, not necessarily a mutant miRNA; only *mir-35* is mutated in the indicated mutant strains. (D) Two-way ANOVA was performed, followed by Dunnett’s multiple comparisons test. Small arrows in (E) indicate positions of *mir-35-41* on ranked x-axis, and arrowhead indicates *mir-35* with mutant variants. (C-E) Mean and SEM of three biological replicates are shown.

Again, we performed miRNA-Taqman qPCR of *mir-35* and its mutant variants, using synthetic miRNAs of known concentrations to generate standard curves for absolute quantification (Figure S1). In embryos, the quantity of *mir-35(mut_3’)* and *mir-35(mir-82_3’)* were increased 20-fold and 139-fold relative to wild type *mir-35*, respectively, while star strand abundances did not reflect these changes (Figure 2C). While potential changes in strand selection or stability of the star strands may confound interpretation of guide:star strand ratio of the mutant duplexes, the large overall increase in the number of molecules deriving from either strand of the 3’ end mutant precursors compared to wild type supports the model that biogenesis-or decay-level effects are contributing to the high abundance of *mir-35(mut_3’)* and *mir-35(mir-82_3’)* in embryos. Despite the caveats to interpreting guide:star ratio of the mutant duplex, we currently favor the model that decay is disrupted over the model that biogenesis is enhanced because the apparent strand specificity of the effect is consistent across both mutant variants. Overall, we postulate that a second regulatory mechanism acts to limit abundance of *mir-35* in the embryo in a manner dependent upon the 3’ end sequence (Figure S4).

We next measured the decay of the *mir-35* 3’ end variants at the embryo to L1 transition. Unlike the seed mutants, the change in the abundance of the *mir-35* 3’ end mutants from the embryo to L1 stage was similar to that of wild type *mir-35* [7-fold for the *mir-35(mut_3’)*, 14-fold for *mir-35(mir-82_3’*), and 8-fold for wild type] (Figure 2B). Likewise *mir-36* was not affected by the mutations (Figure 2B). Again, we confirmed and extended these findings using deep sequencing. Deep sequencing confirmed that the 3’ end variants showed a similar depletion from embryo to L1 stage as wild type *mir-35*, and that no other miRNAs in the *mir-35* family or otherwise were affected (Figure 2D-E, Figure S5, Tables S3 and S4). Thus, the sequence of the 3’ end of the miRNA outside the seed did not affect the decay at this developmental transition.

Overall, we observed that seed mutations do not generally impact embryonic *mir-35* abundance but strongly inhibit its decay at the embryo to L1 transition, whereas 3’ end mutations strongly impact embryonic abundance of *mir-35* but do not affect its decay at the embryo to L1 transition. Taken together, we propose that two mechanisms regulate *mir-35* abundance: a 3’ end-dependent mechanism limits abundance in embryos, while a seed-dependent mechanism drives decay at the transition to L1 (Figure S4A). Given that all positions 3’ of the seed sequence are mutated in the 3’ end mutants, the seed sequence of *mir-35* is not only necessary but also largely sufficient to drive its selective decay at the embryo to L1 transition. Notably, this working model assumes that the 3’ end mutant variants are decayed by the same mechanism as wild type *mir-35* at the embryo to L1 transition; alternatively, if the 3’ end mutant variants are decayed by a novel mechanism, then the 3’ end sequence could still play a role in the embryo to L1 decay of wild type *mir-35*.

### EBAX-1 regulates *mir-35* family abundance in embryos and at the embryo to L1 transition

Given the model that the *mir-35* family is regulated in two phases (Figure S4A), we wondered whether EBAX-1 regulates *mir-35* abundance in either of these developmental windows. To this end, we assayed wild type and *ebax-1(null)* animals. We performed both qPCR in bulk embryos and L1s (similar to other experiments thus far) as well as deep sequencing in hand-picked staged embryos and L1 larvae.

Both assays showed a modest upregulation of *mir-35* family members in embryos: a 1.4-and 1.3-fold increase was observed in *mir-35* and *mir-36*, respectively, by bulk sample qPCR (Figure S4B). These changes were reflected in the star strands as well, which were each increased 1.4-fold (Figure S4C). The deep sequencing data of staged embryo samples also showed a modest increase in embryos, with the greatest change in the comma stage, where *mir-35-41* were upregulated 2.3-fold on average (Figure S4D, Table S5). (Data for star strands was sparse and noisy in this deep sequencing, and therefore likely unreliable to interpret.) Together, these data suggest that EBAX-1 has a modest effect in regulating *mir-35* family abundance in the embryo, and that its effect may impact transcription or biogenesis.

At the embryo to L1 transition, qPCR of bulk samples showed stark stabilization of *mir-35* and *mir-36* in *ebax-1(null)*; decay was 10-and 12-fold for *mir-35* and *mir-36* in wild type, whereas no measurable decay was observed in *ebax-1(null)* (Figure S4B). Deep sequencing results corroborated the impact of *ebax-1(null)* on decay at the embryo to L1 transition; however, the amplitude of this effect was slightly inconsistent. For instance, *mir-35* and *mir-36* were decayed 17-and 8-fold from the comma embryo stage to L1 in wild type, and the amplitude of the decay was reduced to 6-and 5-fold, respectively, from comma to L1 in *ebax-1(null)* (Figure S4D). These discrepancies may arise from the major differences in the two methodologies used, including sample collection and normalization. Unlike for bulk samples, the staged deep sequencing samples were normalized to spike-ins which were added on a per-animal basis, so one possible source of variability is differential lysis efficiency between embryos and L1 stage animals. Nonetheless, both experiments support a role for EBAX-1 in *mir-35* family decay at the embryo to L1 transition.

### *mir-35* variants are tailed and trimmed similarly to wild type *mir-35*

While the seed-dependence and ZSWIM8/EBAX-1-dependence of *mir-35* regulation suggest a TDMD-like mechanism, the dispensability of the 3’ end for developmentally timed decay suggests an alternative mechanism. TDMD is often accompanied by a high prevalence of tailing and trimming during the decay process, which is thought to be due to conformational changes induced by extensive base pairing that expose the 3’ end of the miRNA to exonucleases and nucleotidyltransferases.

To determine whether the *mir-35* family bears this symptom of the TDMD conformation, we examined the prevalence of tailing and trimming. We first examined the level of background in tailing measurements in our experimental and computational pipeline. To this end, synthetic miRNAs were spiked into total RNA after purification, and the amount of tailing called on these miRNAs is considered background since these miRNAs were never present in the context of cellular lysate, so any apparent “tailing” must derive from errors introduced in cloning or sequencing. Tailing was below 1.5% in 317 out of 324 (98%) such spike-in measurements, so tailing below 1.5% is considered background in these datasets. This threshold is marked by a dashed line on all tailing plots.

In embryos and L1s, we observed that miRNAs are tailed to various extents, though generally not very high levels (Figure 3A, Figure S6A). Tailing was mostly mono-uridylation, with some miRNAs displaying significant adenylation or cytidylation, as previously observed (Figure 3A, Figure S6A) (Vieux et al., 2021). Overall tailing and miRNA abundance were not correlated, and the *mir-35* family members were generally high in abundance, with a wide range of tailing frequencies observed across different members (Figure 3B, Table S4).

**Figure 3.**
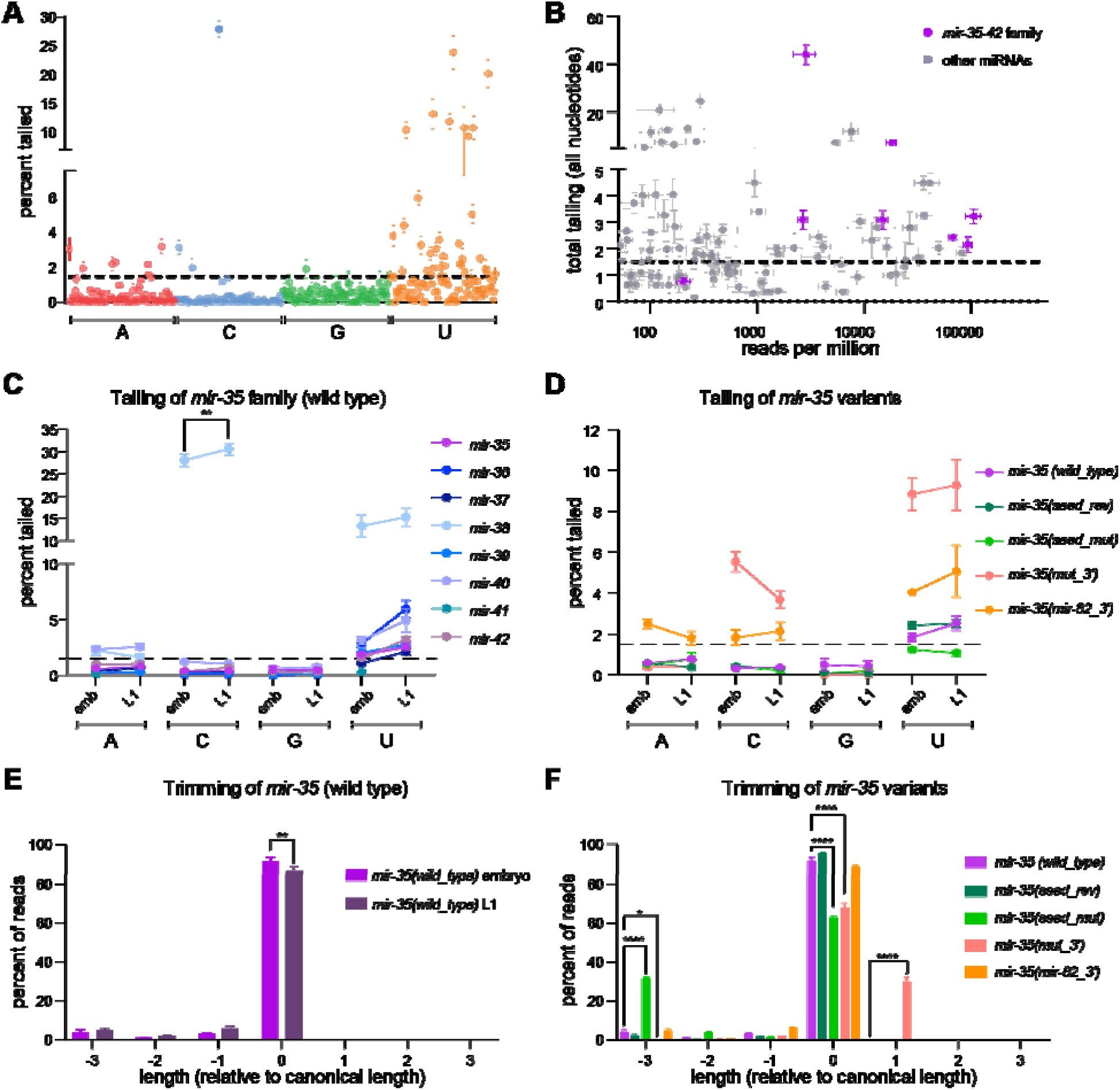
Changes in tailing and trimming of *mir-35* variants do not correlate with changes in decay. (A) Percent of reads with single nucleotide addition to the 3’ end is shown for each miRNA with >50 RPM in embryo. (B) Total tailing (sum of all single nucleotide tails) versus abundance (RPM) is shown for all miRNAs with >50 RPM in embryo. The *mir-35* family is highlighted in purple. (C-D) Percent of reads with single nucleotide additions to the *mir-35* family (C) and the *mir-35* variants (D) in the embryo and L1. (E-F) Percent of reads of each length (excluding tail), relative to the canonical length of *mir-35* in wild type embryo and L1 (E) or *mir-35* variants in embryo (F). (A-F) Mean and SEM are shown for six biological replicates for wild type and three biological replicates for all *mir-35* variant strains. (C-F) For each nucleotide, one-way ANOVA was performed, followed by Sidak’s multiple comparison test. Significant relationships in panel D are described in the text, but not indicated on the graph for simplicity. *p-value < 0.05, **p-value < 0.01, ****p-value < 0.0001.

We and others previously observed slight increases in the prevalence of tailed and trimmed miRNAs as miRNAs approach decay (Baccarini et al., 2011; Kingston and Bartel, 2019; Vieux et al., 2021). In TDMD, miRNAs often experience very high levels of tailing and/or trimming (generally ≥20-40% tailed or trimmed isoforms) (Ameres et al., 2010; Baccarini et al., 2011; Bitetti et al., 2018; Cazalla et al., 2010; Ghini et al., 2018; Kleaveland et al., 2018; Li et al., 2021; Marcinowski et al., 2012). We hypothesized that the prevalence of tailed isoforms might increase at the embryo to L1 transition as the *mir-35* family members undergo decay. Small increases in miRNA tailing and trimming were observed, but in most cases, these were not statistically significant, and the prevalence of modified isoforms remained modest (Figure 3C, 3E, Table S4).

We next examined the prevalence of tailed isoforms in the context of mutant versions of *mir-35*. Significant changes in tailing were observed, but interestingly, these did not correlate with changes in rates of decay (Figure 3D, Table S4). For instance, *mir-35(mut_3’)* was significantly more cytidylated and uridylated than wild type *mir-35* in embryo and L1, and *mir-35(mir-82_3’)* was significantly more adenylated, cytidylated, and uridylated than wild type *mir-35* in both stages (p-value < 0.05 for all aforementioned comparisons) (Figure 3D). However, these two *mir-35* variants displayed decay similar to that of wild type *mir-35* at the embryo to L1 transition (Figure 2B). In contrast, *mir-35(seed_rev)* and *mir-35(seed_mut)* show similar amounts of tailing to the wild type *mir-35* (Figure 3D), despite these variants’ dramatically altered decay at the embryo to L1 transition (Figure 1C). Oligonucleotide tails were much less frequent than single nucleotide tails, with di-nucleotide tails occurring about an order of magnitude less frequently than the corresponding single nucleotide tail; again *mir-35(seed_rev)* showed a very similar complement of oligo-nucleotide tails to wild type *mir-35* (Table S6). Thus, changes in tailing did not correlate with changes in decay.

We next examined trimming of *mir-35* variants. Like tailing, trimming varied widely among *mir-35* variants, but not in a manner that correlated with the rate of decay. For instance, trimming increased most for *mir-35(seed_mut)*, despite the enhanced stability of this variant (Figure 3F, Figure S6B). In contrast, *mir-35(seed_rev)* – which shows similarly enhanced stability – had no significant change in trimming (Figure 3F, Figure S6B). This isoform analysis also showed that *mir-35(mut_3’)* yields two major isoforms from biogenesis, the canonical 22-nt isoform and a 23-nt isoform which is extended by 1nt at the 3’ end (Figure 3F, Figure S6B). Analysis of deep sequencing data showed that both isoforms are decayed similarly at the embryo to L1 transition (Figure S7). Overall, changes in trimming did not correlate with changes in decay.

All together, these data show that the tailing and trimming of the *mir-35* family are much lower in prevalence than in most known instances of TDMD, and that the incidence of trimmed and tailed isoforms across *mir-35* variants did not correlate with rate of decay at the embryo to L1 transition. Together with the dispensability of the 3’ end sequences of *mir-35* for decay, this suggests that the mechanism of decay of *mir-35* differs from previously described examples of TDMD.

### Reintroducing miRNA-target interactions does not restore decay of seed mutant variants of *mir-35*

To further investigate the mechanism of *mir-35* family decay at the embryo to L1 transition, we examined the possible involvement of complementary RNA molecules as in TDMD. Decay of *mir-35* at the embryo to L1 transition is dependent on its seed sequence but not its 3’ end sequences, and canonical targets were previously shown to regulate miRNA stability in *C. elegans* (Chatterjee et al., 2011). We therefore wondered whether canonical miRNA:target interactions (rather than TDMD-like base-pairing) might play a role in mediating this decay. The *mir-35(seed_rev)* variant is predicted to have fewer target molecules expressed in embryos compared to those of wild type *mir-35*: the *mir-35(seed_rev)* target pool is 59% that of wild type based on target prediction and relative expression according to RNAseq (Agarwal et al., 2015; Grün et al., 2014). The lower dose of canonical target interactions may influence decay, or alternatively, wild type *mir-35* targets may have unknown properties required for decay. To test these hypotheses, we restored canonical target interactions for *mir-35(seed_rev)* to determine whether this restored developmentally-timed decay.

To this end, we sought to alter a similar stoichiometric proportion of the pool of *mir-35* family miRNAs and the pool of *mir-35* family targets. *mir-35* makes up 20% of the *mir-35-42* miRNA molecules in embryos based on quantitative deep sequencing (Dexheimer et al., 2020), so we selected three target genes that together also make up 20% of the target molecules (as estimated from embryo RNAseq datasets) (Grün et al., 2014). These genes – *egl-1, nhl-2*, and *sup-26 –* were also selected because they are all validated targets that influence physiology downstream of *mir-35-42* (Kagias and Pocock, 2015; McJunkin and Ambros, 2017; Sherrard et al., 2017; Tran et al., 2019; Wu et al., 2010; Yang et al., 2020b). Using CRISPR, we made mutations to the *mir-35* family binding site in the 3’UTR of these three target genes. These mutations enable binding by *mir-35(seed_rev)* rather than wild type *mir-35*, and we have previously shown that *mir-35(seed_rev)* represses such targets (Figure 4A) (Yang et al., 2020b).

**Figure 4.**
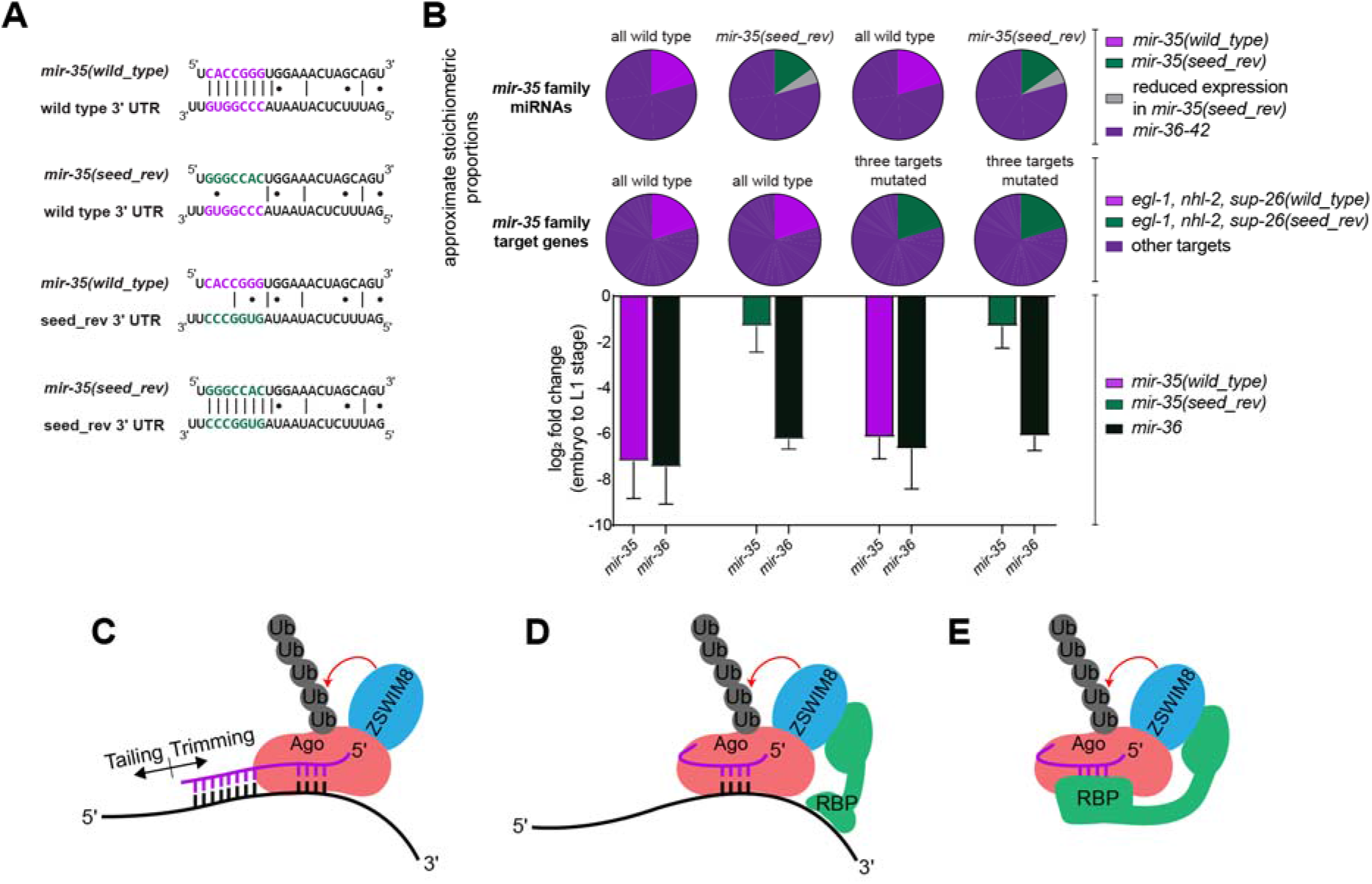
Reintroducing miRNA-target interactions does not restore decay of a seed mutant variant of *mir-35*. (A) Target binding sites for *mir-35(seed_rev)* were introduced to the 3’UTR of three known *mir-35* targets (*egl-1, sup-26* and *nhl-2*), either in a *mir-35(wild_type)* or a *mir-35(seed_rev)* background. An example of the relevant miRNA-target interactions at the *egl-1* 3’UTR are shown. (B) Top: Pie charts represent an estimate of the proportion of the *mir-35* family molecules and the *mir-35* family target molecules that are mutated in each strain. Bottom: Log_2_(fold change) from embryo to L1 in the indicated strains, as measured by Taqman qPCR. Mean and SEM of three biological replicates are shown. (C) Model of conventional TDMD. (D-E) Alternative models for regulation of *mir-35* family decay, in which the seed sequence is recognized by a complementary RNA (D) or an RNA-binding protein (RBP) (E).

We performed qPCR to measure the miRNA levels in the embryo and L1 stages. Again, we observed a significant reduction in wild type *mir-35* from embryo to L1 and attenuated decay of the *mir-35(seed_rev)* (Figure 4B). Wild type *mir-35* decay was not affected by the mutations of the target sites (Figure 4B). When *mir-35(seed_rev)* was combined with the mutant targets containing complementary binding sites, decay was similar to *mir-35(seed_rev)* without engineered target interactions (Figure 4B). Thus, restoring interactions with canonical target genes is not sufficient to restore turnover of the *mir-35* seed mutant.

## Discussion

Here we investigate the regulation of the embryonically-expressed *mir-35* family during development. We show that the decay of these miRNAs at the embryo to L1 transition is regulated post-transcriptionally, since mutations in the seed sequence of *mir-35* decouple its regulation from that of its clustermates on the same transcript, strongly supporting a selective decay mechanism.

The seed sequence of *mir-35* is not only necessary for this selective decay, but is also largely sufficient since mutations in the 3’ end of the miRNA do not disrupt decay at the embryo to L1 transition. This model assumes that the 3’ end mutant variants are decayed by the same mechanism as wild type *mir-35* in this developmental window, and future mechanistic studies will test that assumption. We also observe that the 3’ end regulates *mir-35* abundance in the embryo, in what may be a decay-level effect. We postulate that, whereas a seed-dependent decay mechanism enacts developmentally-timed decay, a 3’ end-dependent mechanism limits *mir-35* abundance in the embryo (Figure S4).

While the TDMD factor ZSWIM8/EBAX-1 regulates *mir-35* abundance, our data argues that the mechanisms of *mir-35* regulation differ from TDMD in key aspects (Figure 4C-E). First, the decay at embryo to L1 does not require the 3’ end sequences which would be involved in base pairing to a typical TDMD trigger RNA. Second, the decay is not accompanied by high levels of tailing or trimming. Furthermore, seed mutations that reduce decay do not reduce tailing or trimming. Together, these data suggest that the *mir-35* family is post-transcriptionally regulated by a novel seed-dependent mechanism. We further observed that the altered regulation of seed mutants was not due to loss of target interactions, since restoring these interactions did not restore decay.

We propose a model for *mir-35* family decay wherein ZSWIM8/EBAX-1 is recruited to degrade the miRISC in a seed-dependent manner that does not require extensive 3’ end base pairing. How the seed is recognized and how ZSWIM8/EBAX-1 is recruited in this process will be an area of ongoing study. Like TDMD, a trigger RNA may base pair with the *mir-35* family seed sequence and recruit an RNA binding protein, which can in turn recruit ZSWIM8/EBAX-1 (Figure 4D). Alternatively, the trigger RNA could induce conformational changes in Argonaute that directly recruit ZSWIM8/EBAX-1, similar to proposed models of TDMD. A third possibility is that no trigger RNA is involved in seed recognition for decay; in this case an RBP could bind the *mir-35* seed to recruit ZSWIM8/EBAX-1 or induce Argonaute conformational changes (Figure 4E). Because of the large number of possible trigger RNAs or trigger RBPs, further elucidating this mechanism will require large scale biochemical and genetic screens. Better understanding the mechanism of *mir-35* family recognition will further test the model of the sufficiency of the seed sequence alone for this selective regulation.

Understanding the seed sequence-specific decay mechanism regulating *mir-35* will have broad impact. Such seed-specific mechanisms are likely to be present in other biological systems because they allow for simultaneous regulation of redundant paralogs, enabling dynamic regulation of a miRNA seed family’s targets. Outside functioning in normal physiology, seed-specific decay mechanisms could be an attractive avenue for modulating abundance of specific miRNA families and their target genes in disease.

## Limitations of the Study

A general limitation to the study is that only *mir-35* was modified and examined; whether the strength of the impact of these mutations is similar in the context of *mir-36-42* will be a matter of future investigation. Having not examined processing intermediates, we have as yet an incomplete understanding of how the mutations introduced in *mir-35* affect its biogenesis. Related to this caveat, we have not directly measured the decay of *mir-35* 3’ end variants in the embryo stage due to technical difficulty, leaving some remaining ambiguity as to the molecular basis of upregulation of *mir-35* 3’ end mutant variants in the embryo. Furthermore, the decay of the 3’ end mutant variants of *mir-35* at the embryo to L1 transition occurs at a similar rate as wild type; our interpretation is that these variants are subject to the same mechanism that targets wild type *mir-35* at the embryo to L1 transition. However, if the 3’ end variants are targeted by a distinct mechanism of decay in the same developmental time window, this could obscure the 3’ end’s role in the regulation of wild type *mir-35*.

## Materials and Methods

### General *C. elegans* culture and maintenance

*C. elegans* were maintained at 20°C on NGM seeded with OP50. For large-scale harvest of embryos, 8,000 starved L1s were plated onto a 10cm plate with a large lawn of OP50. The worms were re-fed with concentrated OP50 48 hours later. At 96 hours after initial plating, the gravid adults were harvested by bleaching to collect large quantities of embryos.

### Liquid culture

For experiments in which bulk sample deep sequencing was performed (Figures 1 and 2), worms were grown in liquid culture as previously described and harvested with some modifications (Zanin et al., 2011). Briefly, the gravid worms were harvested by centrifugation at 3000xg for 2 minutes in 50mL conical tubes. They were washed once with room temperature water and then pelleted. The volume of the sample was brought up to 28mL with water, and then 4mL of 5M NaOH and 8mL of 4% sodium hypochlorite were added. The tubes were immediately shaken vigorously for 2 minutes and allowed to rest on the bench for 1 minute, and this shaking and resting was repeated three times. The worms were immediately centrifuged at 3000xg for 2 minutes. The supernatant was decanted, and the worms were washed four times with 45mL of water. The synchronized embryos were either collected for the embryo samples, or M9 with cholesterol was added, and the worms were placed on a rocker at 20°C overnight to obtain a population of synchronized, starved L1 worms.

### CRISPR/Cas9-mediated genome editing

For all CRISPR experiments, pre-assembled Cas9 RNPs were injected into germlines along with short homology-directed repair templates with ∼35-nt homology arms (Paix et al., 2014). For all CRISPR injections, one of the guide RNAs used targeted *dpy-10* as a visible marker to select plates with efficient genome editing (Arribere et al., 2014). crRNAs and tracrRNA were ordered from IDT (Alt-R) or Dharmacon (Edit-R), and annealed at 10µM in IDT duplex buffer by heating to 95°C for 5 minutes and then cooling to room temperature. Injection mixes contained 2-4µM Cas9, 4µM total of pre-annealed gRNAs (comprised of gRNAs targeting *dpy-10* and the site of interest), 0.8µM of the *dpy-10* donor oligonucleotide, and the homology directed donor at 40-100ng/µl (Table S2).

Mutations to *mir-35* were made by two rounds of CRISPR. First, as previously described (Yang et al., 2020b), two gRNAs recognizing the protospacers TTTCCATTAGAACTATCACC and ATTGCTGGTTTCTTCCACAG were used to create a 50bp deletion at the *mir-35* locus. This allele is *mir-35(cdb2):*

GCTGGTTTCTTCCACAGT-50bp_del-CTTTTCCACTTGCTCCAC. The strain carrying *mir-35(cdb2)* was then injected with homology-directed repair donors, along with a gRNA (GGAGCAAGTGGAAAAGACTG) recognizing a sequence which is created by the *mir-35(cdb2)* mutation. See Table S1 for allele and strain details and Table S2 for all donor oligonucleotides.

### Deep sequencing library preparation and data analysis

Library preparation was performed using the NEBNext Small RNA Library Prep Set for Illumina with modifications as previously described (Vieux et al., 2021). Briefly, size selection was performed only after reverse transcription, using 8% urea gels to purify ∼65-75nt RT products. Prior to loading on the gel, each RT reaction was treated with 5000units of RNAse H (New England Biolabs) for 30 minutes at 37°C. For bulk embryo and L1 samples, 15 PCR cycles were performed. For samples of 20 staged embryos or 20 corresponding L1s, 15-20 PCR cycles were performed. Sequence analysis was performed on the NIH High Performance Computing Cluster. The 3’ adapter sequence was trimmed using Cutadapt 3.4 (Martin, 2011). The reads were mapped to a custom genome file which was comprised of *C. elegans* genome WS280 with an additional chromosome containing the sequences of the spike-in miRNAs and the mutant *mir-35* precursors with flanking genomic sequence. Mapping was performed using bowtie2 2.4.4 (Langmead et al., 2009) with the following settings: --no-unal --end-to-end --sensitive. BAM files were sorted and indexed using samtools 1.13 (Danecek et al., 2021). Reads were assigned to miRNAs using htseq 0.13.5 (Anders et al., 2015) with the following settings: --mode union -- nonunique fraction -a 0. The htseq analysis was performed using a gff file modified from mirGeneDB (Fromm et al., 2015) by replacing mirGeneDB IDs with miRbase IDs (Kozomara and Griffiths-Jones, 2014) and adding the intervals corresponding to the spike-in miRNAs and the *mir-35* mutant miRNAs in the custom genome file. For analysis of tailing and trimming, the Tailor package (Chou et al., 2015) was used with the genome file described above and FASTA files derived from mirGeneDB, but with IDs replaced by miRbase IDs and sequences for spike-in miRNAs and the *mir-35* mutant miRNAs appended.

To assess the stoichiometry of the potential targets of *mir-35(seed_rev)*, TargetScan 7.0 was used to predict binding sites for *mir-35(seed_rev)*. Expression data from (Grün et al., 2014) was used to infer relative expression of predicted target genes.

### RNA isolation

Total RNA was isolated from bulk samples by resuspending the sample in the recommended volume of Trizol reagent (Life Technologies), followed by vortexing at room temperature for 15 minutes, followed by preparation according to the Trizol manufacturer’s instructions. After preparation, ten spike-in oligos (see Table S2) were added at a final concentration of 1pg/µl each in 100ng/µl total RNA prior to deep sequencing library preparation.

For staged embryo samples, 20 staged embryos or 20 L1s (synchronized by starvation for 24h) were collected by hand. Samples were snap frozen in Trizol LS reagent. Prior to purification, 0.9pg of each spike-in oligo (Table S2) was added to each sample. Trizol LS-resuspended samples were subjected to three freeze-thaw cycles to promote lysis and then vortexed for 15 minutes at room temperature prior to purification according to the Trizol LS manufacturer’s instructions.

### Taqman miRNA qPCR

For all miRNA qPCR, 5µl reverse transcription reactions were performed using the TaqMan MicroRNA Reverse Transcription kit (ThermoFisher). For all samples, 1.66µl of total RNA at 6ng/µl was used in the reverse transcription. RT reactions were diluted 1:4 and 1.33µl was used in a 5µl qPCR reaction prepared using Taqman miRNA probes with the Taqman Universal Mastermix II with UNG (ThermoFisher). Reactions were run in triplicate on the Applied Biosystems QuantStudio Pro 6.

## Data Availability

All raw sequence data has been deposited in NCBI Sequence Read Archive (SRA) under accession number PRJNA782102.

## Acknowledgments

We thank WormBase, the NIDDK Genomics Core, the NCI Genomics Core, and NIH High Performance Computing. Strains are regularly received from the CGC, which is funded by the NIH Office of Research Infrastructure Programs (P40 OD010440). Thank you to Yishi Jin for the *ebax-1(null)* strain. Thank you to members of the McJunkin lab, Eric Miska, Kenneth Murfitt, Michael Lichten, Joana Vidigal, John Kim, and Leemor Joshua-Tor for helpful discussions.

## References

Agarwal, V., Bell, G.W., Nam, J.W., and Bartel, D.P. (2015). Predicting effective microRNA target sites in mammalian mRNAs. Elife 4. https://doi.org/10.7554/ELIFE.05005.

Alvarez-Saavedra, E., and Horvitz, H.R. (2010). Many families of C. elegans microRNAs are not essential for development or viability. Curr. Biol. 20, 367–373. https://doi.org/10.1016/j.cub.2009.12.051.

Ameres, S.L., Horwich, M.D., Hung, J.-H., Xu, J., Ghildiyal, M., Weng, Z., and Zamore, P.D. (2010). Target RNA-directed trimming and tailing of small silencing RNAs. Science 328, 1534–1539. https://doi.org/10.1126/science.1187058.

Anders, S., Pyl, P.T., and Huber, W. (2015). HTSeq—a Python framework to work with high-throughput sequencing data. Bioinformatics 31, 166–169. https://doi.org/10.1093/BIOINFORMATICS/BTU638.

Arribere, J.A., Bell, R.T., Fu, B.X.H., Artiles, K.L., Hartman, P.S., and Fire, A.Z. (2014). Efficient marker-free recovery of custom genetic modifications with CRISPR/Cas9 in Caenorhabditis elegans. Genetics 198, 837–846. https://doi.org/10.1534/genetics.114.169730.

Baccarini, A., Chauhan, H., Gardner, T.J., Jayaprakash, A.D., Sachidanandam, R., and Brown, B.D. (2011). Kinetic analysis reveals the fate of a microRNA following target regulation in mammalian cells. Curr. Biol. 21, 369–376. https://doi.org/10.1016/j.cub.2011.01.067.

Bail, S., Swerdel, M., Liu, H., Jiao, X., Goff, L.A., Hart, R.P., and Kiledjian, M. (2010). Differential regulation of microRNA stability. RNA 16, 1032–1039. https://doi.org/10.1261/rna.1851510.

Bernstein, E., Caudy, A.A., Hammond, S.M., and Hannon, G.J. (2001). Role for a bidentate ribonuclease in the initiation step of RNA interference. Nature 409, 363–366. https://doi.org/10.1038/35053110.

Bitetti, A., Mallory, A.C., Golini, E., Carrieri, C., Carreño Gutiérrez, H., Perlas, E., Pérez-Rico, Y.A., Tocchini-Valentini, G.P., Enright, A.J., Norton, W.H.J., et al. (2018). MicroRNA degradation by a conserved target RNA regulates animal behavior. Nat. Struct. Mol. Biol. 25, 244–251. https://doi.org/10.1038/s41594-018-0032-x.

Boele, J., Persson, H., Shin, J.W., Ishizu, Y., Newie, I.S., Sokilde, R., Hawkins, S.M., Coarfa, C., Ikeda, K., Takayama, K. -i., et al. (2014). PAPD5-mediated 3’ adenylation and subsequent degradation of miR-21 is disrupted in proliferative disease. Proc. Natl. Acad. Sci. 111, 11467–11472. https://doi.org/10.1073/pnas.1317751111.

Bossé, G.D., Rüegger, S., Ow, M.C., Vasquez-Rifo, A., Rondeau, E.L., Ambros, V.R., and Simard, M.J. (2013). The decapping scavenger enzyme DCS-1 controls microRNA levels in Caenorhabditis elegans. Mol. Cell 50, 281–287. https://doi.org/10.1016/j.molcel.2013.02.023.

Brancati, G., and Großhans, H. (2018). An interplay of miRNA abundance and target site architecture determines miRNA activity and specificity. Nucleic Acids Res. 46, 3259–3269. https://doi.org/10.1093/nar/gky201.

Brennecke, J., Stark, A., Russell, R.B., and Cohen, S.M. (2005). Principles of microRNA-target recognition. PLoS Biol. 3, e85. https://doi.org/10.1371/journal.pbio.0030085.

Broughton, J.P., Lovci, M.T., Huang, J.L., Yeo, G.W., and Pasquinelli, A.E. (2016). Pairing beyond the Seed Supports MicroRNA Targeting Specificity. Mol. Cell 64, 320–333. https://doi.org/10.1016/j.molcel.2016.09.004.

Cazalla, D., Yario, T., Steitz, J.A., and Steitz, J. (2010). Down-regulation of a host microRNA by a Herpesvirus saimiri noncoding RNA. Science 328, 1563–1566. https://doi.org/10.1126/science.1187197.

Chatterjee, S., Grosshans, H., and Großhans, H. (2009). Active turnover modulates mature microRNA activity in Caenorhabditis elegans. Nature 461, 546–549. https://doi.org/10.1038/nature08349.

Chatterjee, S., Fasler, M., Büssing, I., and Grosshans, H. (2011). Target-mediated protection of endogenous microRNAs in C. elegans. Dev. Cell 20, 388–396. https://doi.org/10.1016/j.devcel.2011.02.008.

Chou, M. Te, Han, B.W., Hsiao, C.P., Zamore, P.D., Weng, Z., and Hung, J.H. (2015). Tailor: a computational framework for detecting non-templated tailing of small silencing RNAs. Nucleic Acids Res. 43. https://doi.org/10.1093/NAR/GKV537.

Dallaire, A., Frédérick, P.-M., and Simard, M.J. (2018). Somatic and Germline MicroRNAs Form Distinct Silencing Complexes to Regulate Their Target mRNAs Differently. Dev. Cell 47, 239–247.e4. https://doi.org/10.1016/j.devcel.2018.08.022.

Danecek, P., Bonfield, J.K., Liddle, J., Marshall, J., Ohan, V., Pollard, M.O., Whitwham, A., Keane, T., McCarthy, S.A., Davies, R.M., et al. (2021). Twelve years of SAMtools and BCFtools. Gigascience 10. https://doi.org/10.1093/GIGASCIENCE/GIAB008.

Denli, A.M., Tops, B.B.J., Plasterk, R.H. a, Ketting, R.F., and Hannon, G.J. (2004). Processing of primary microRNAs by the Microprocessor complex. Nature 432, 231–235. https://doi.org/10.1038/nature03049.

Dexheimer, P.J., and Cochella, L. (2020). MicroRNAs: From Mechanism to Organism. Front. Cell Dev. Biol. 8, 409. https://doi.org/10.3389/fcell.2020.00409.

Dexheimer, P.J., Wang, J., and Cochella, L. (2020). Two microRNAs are sufficient for embryogenesis in C. elegans. BioRxiv 2020.06.28.176024. https://doi.org/10.1101/2020.06.28.176024.

Doll, M.A., Soltanmohammadi, N., and Schumacher, B. (2019). ALG-2/AGO-Dependent mir-35 Family Regulates DNA Damage-Induced Apoptosis Through MPK-1/ERK MAPK Signaling Downstream of the Core Apoptotic Machinery in Caenorhabditis elegans. Genetics 213, 173–194. https://doi.org/10.1534/GENETICS.119.302458.

Elbarbary, R.A., Miyoshi, K., Myers, J.R., Du, P., Ashton, J.M., Tian, B., and Maquat, L.E. (2017a). Tudor-SN-mediated endonucleolytic decay of human cell microRNAs promotes G1/S phase transition. Science 356, 859–862. https://doi.org/10.1126/science.aai9372.

Elbarbary, R.A., Miyoshi, K., Hedaya, O., Myers, J.R., and Maquat, L.E. (2017b). UPF1 helicase promotes TSN-mediated miRNA decay. Genes Dev. 31, 1483–1493. https://doi.org/10.1101/gad.303537.117.

Fang, W., and Bartel, D.P. (2015). The Menu of Features that Define Primary MicroRNAs and Enable De Novo Design of MicroRNA Genes. Mol. Cell 60, 131–145. https://doi.org/10.1016/j.molcel.2015.08.015.

Flamand, M.N., Wu, E., Vashisht, A., Jannot, G., Keiper, B.D., Simard, M.J., Wohlschlegel, J., and Duchaine, T.F. (2016). Poly(A)-binding proteins are required for microRNA-mediated silencing and to promote target deadenylation in C. elegans. Nucleic Acids Res. 44, 5924–5935. https://doi.org/10.1093/NAR/GKW276.

Flamand, M.N., Gan, H.H., Mayya, V.K., Gunsalus, K.C., and Duchaine, T.F. (2017). A non-canonical site reveals the cooperative mechanisms of microRNA-mediated silencing. Nucleic Acids Res. 1–14. https://doi.org/10.1093/nar/gkx340.

Fromm, B., Billipp, T., Peck, L.E., Johansen, M., Tarver, J.E., King, B.L., Newcomb, J.M., Sempere, L.F., Flatmark, K., Hovig, E., et al. (2015). A Uniform System for the Annotation of Vertebrate microRNA Genes and the Evolution of the Human microRNAome. http://Dx.Doi.Org/10.1146/Annurev-Genet-120213-092023 49, 213–242. https://doi.org/10.1146/ANNUREV-GENET-120213-092023.

Ghini, F., Rubolino, C., Climent, M., Simeone, I., Marzi, M.J., and Nicassio, F. (2018). Endogenous transcripts control miRNA levels and activity in mammalian cells by target-directed miRNA degradation. Nat. Commun. 9, 3119. https://doi.org/10.1038/s41467-018-05182-9.

Gregory, R.I., Yan, K.-P., Amuthan, G., Chendrimada, T., Doratotaj, B., Cooch, N., and Shiekhattar, R. (2004). The Microprocessor complex mediates the genesis of microRNAs. Nature 432, 235–240. https://doi.org/10.1038/nature03120.

Grishok, A., Pasquinelli, A.E., Conte, D., Li, N., Parrish, S., Ha, I., Baillie, D.L., Fire, A., Ruvkun, G., and Mello, C.C. (2001). Genes and mechanisms related to RNA interference regulate expression of the small temporal RNAs that control C. elegans developmental timing. Cell 106, 23–34..

Grün, D., Kirchner, M., Thierfelder, N., Stoeckius, M., Selbach, M., and Rajewsky, N. (2014). Conservation of mRNA and protein expression during development of C. elegans. Cell Rep. 6, 565–577. https://doi.org/10.1016/J.CELREP.2014.01.001.

Han, J., Lee, Y., Yeom, K.-H., Kim, Y.-K., Jin, H., and Kim, V.N. (2004). The Drosha-DGCR8 complex in primary microRNA processing. Genes Dev. 18, 3016–3027. https://doi.org/10.1101/gad.1262504.

Han, J., Lee, Y., Yeom, K.-H., Nam, J.-W., Heo, I., Rhee, J.-K., Sohn, S.Y., Cho, Y., Zhang, B.-T., and Kim, V.N. (2006). Molecular basis for the recognition of primary microRNAs by the Drosha-DGCR8 complex. Cell 125, 887–901. https://doi.org/10.1016/j.cell.2006.03.043.

Han, J., LaVigne, C.A., Jones, B.T., Zhang, H., Gillett, F., and Mendell, J.T. (2020). A ubiquitin ligase mediates target-directed microRNA decay independently of tailing and trimming. Science 370, eabc9546. https://doi.org/10.1126/science.abc9546.

Helwak, A., Kudla, G., Dudnakova, T., and Tollervey, D. (2013). Mapping the Human miRNA Interactome by CLASH Reveals Frequent Noncanonical Binding. Cell 153, 654. https://doi.org/10.1016/J.CELL.2013.03.043.

Hutvágner, G., McLachlan, J., Pasquinelli, A.E., Bálint, E., Tuschl, T., and Zamore, P.D. (2001). A cellular function for the RNA-interference enzyme Dicer in the maturation of the let-7 small temporal RNA. Science 293, 834–838. https://doi.org/10.1126/science.1062961.

Iwasaki, S., Kobayashi, M., Yoda, M., Sakaguchi, Y., Katsuma, S., Suzuki, T., and Tomari, Y. (2010). Hsc70/Hsp90 chaperone machinery mediates ATP-dependent RISC loading of small RNA duplexes. Mol. Cell 39, 292–299. https://doi.org/10.1016/j.molcel.2010.05.015.

Iwasaki, S., Sasaki, H.M., Sakaguchi, Y., Suzuki, T., Tadakuma, H., and Tomari, Y. (2015). Defining fundamental steps in the assembly of the Drosophila RNAi enzyme complex. Nature 521, 533–536. https://doi.org/10.1038/nature14254.

Kagias, K., and Pocock, R. (2015). microRNA regulation of the embryonic hypoxic response in Caenorhabditis elegans. Sci. Rep. 5, 11284. https://doi.org/10.1038/srep11284.

Kato, M., de Lencastre, A., Pincus, Z., and Slack, F.J. (2009). Dynamic expression of small non-coding RNAs, including novel microRNAs and piRNAs/21U-RNAs, during Caenorhabditis elegans development. Genome Biol. 10, R54. https://doi.org/10.1186/gb-2009-10-5-r54.

Katoh, T., Hojo, H., and Suzuki, T. (2015). Destabilization of microRNAs in human cells by 3′ deadenylation mediated by PARN and CUGBP1. Nucleic Acids Res. 43, 7521–7534. https://doi.org/10.1093/nar/gkv669.

Ketting, R.F., Fischer, S.E., Bernstein, E., Sijen, T., Hannon, G.J., and Plasterk, R.H. (2001). Dicer functions in RNA interference and in synthesis of small RNA involved in developmental timing in C. elegans. Genes Dev. 15, 2654–2659. https://doi.org/10.1101/gad.927801.

Kingston, E.R., and Bartel, D.P. (2019). Global analyses of the dynamics of mammalian microRNA metabolism. Genome Res. 29, 1777–1790. https://doi.org/10.1101/gr.251421.119.

Kleaveland, B., Shi, C.Y., Stefano, J., and Bartel, D.P. (2018). A Network of Noncoding Regulatory RNAs Acts in the Mammalian Brain. Cell 174, 350–362.e17. https://doi.org/10.1016/j.cell.2018.05.022.

Knight, S.W., and Bass, B.L. (2001). A role for the RNase III enzyme DCR-1 in RNA interference and germ line development in Caenorhabditis elegans. Science 293, 2269–2271. https://doi.org/10.1126/science.1062039.

Knouf, E.C., Wyman, S.K., and Tewari, M. (2013). The human TUT1 nucleotidyl transferase as a global regulator of microRNA abundance. PLoS One 8, e69630. https://doi.org/10.1371/journal.pone.0069630.

Kozomara, A., and Griffiths-Jones, S. (2014). MiRBase: Annotating high confidence microRNAs using deep sequencing data. Nucleic Acids Res. 42, D68–D73. https://doi.org/10.1093/nar/gkt1181.

Krol, J., Busskamp, V., Markiewicz, I., Stadler, M.B., Ribi, S., Richter, J., Duebel, J., Bicker, S., Fehling, H.J., Schübeler, D., et al. (2010). Characterizing light-regulated retinal microRNAs reveals rapid turnover as a common property of neuronal microRNAs. Cell 141, 618–631. https://doi.org/10.1016/j.cell.2010.03.039.

Landthaler, M., Yalcin, A., and Tuschl, T. (2004). The Human DiGeorge Syndrome Critical Region Gene 8 and Its D. melanogaster Homolog Are Required for miRNA Biogenesis. Curr. Biol. 14, 2162–2167. https://doi.org/10.1016/j.cub.2004.11.001.

Langmead, B., Trapnell, C., Pop, M., and Salzberg, S.L. (2009). Ultrafast and memory-efficient alignment of short DNA sequences to the human genome. Genome Biol. 10, R25. https://doi.org/10.1186/gb-2009-10-3-r25.

Lee, D., Park, D., Park, J.H., Kim, J.H., and Shin, C. (2019). Poly(A)-specific ribonuclease sculpts the 3′ ends of microRNAs. RNA 25, 388–405. https://doi.org/10.1261/rna.069633.118.

Lee, M., Choi, Y., Kim, K., Jin, H., Lim, J., Nguyen, T.A.A., Yang, J., Jeong, M., Giraldez, A.J.J., Yang, H., et al. (2014). Adenylation of maternally inherited microRNAs by Wispy. Mol. Cell 56, 696–707. https://doi.org/10.1016/j.molcel.2014.10.011.

Lehrbach, N.J., Castro, C., Murfitt, K.J., Abreu-Goodger, C., Griffin, J.L., and Miska, E.A. (2012). Post-developmental microRNA expression is required for normal physiology, and regulates aging in parallel to insulin/IGF-1 signaling in C. elegans. RNA 18, 2220–2235. https://doi.org/10.1261/rna.035402.112.

Lewis, B.P., Shih, I., Jones-Rhoades, M.W., Bartel, D.P., and Burge, C.B. (2003). Prediction of mammalian microRNA targets. Cell 115, 787–798. https://doi.org/10.1016/s0092-8674(03)01018-3.

Li, L., Sheng, P., Li, T., Fields, C.J., Hiers, N.M., Wang, Y., Li, J., Guardia, C.M., Licht, J.D., and Xie, M. (2021). Widespread microRNA degradation elements in target mRNAs can assist the encoded proteins. Genes Dev. 35, 1595–1609. https://doi.org/10.1101/GAD.348874.121/-/DC1.

Libri, V., Helwak, A., Miesen, P., Santhakumar, D., Borger, J.G., Kudla, G., Grey, F., Tollervey, D., and Buck, A.H. (2012). Murine cytomegalovirus encodes a miR-27 inhibitor disguised as a target. Proc. Natl. Acad. Sci. U. S. A. 109, 279–284. https://doi.org/10.1073/pnas.1114204109.

Liu, M., Liu, P., Zhang, L., Cai, Q., Gao, G., Zhang, W., Zhu, Z., Liu, D., and Fan, Q. (2011). mir-35 is involved in intestine cell G1/S transition and germ cell proliferation in C. elegans. Cell Res. 21, 1605–1618. https://doi.org/10.1038/cr.2011.102.

Ma, H., Wu, Y., Choi, J.-G., and Wu, H. (2013). Lower and upper stem-single-stranded RNA junctions together determine the Drosha cleavage site. Proc. Natl. Acad. Sci. U. S. A. 110, 20687–20692. https://doi.org/10.1073/pnas.1311639110.

Marcinowski, L., Tanguy, M., Krmpotic, A., Rädle, B., Lisnić, V.J., Tuddenham, L., Chane-Woon-Ming, B., Ruzsics, Z., Erhard, F., Benkartek, C., et al. (2012). Degradation of cellular mir-27 by a novel, highly abundant viral transcript is important for efficient virus replication in vivo. PLoS Pathog. 8, e1002510. https://doi.org/10.1371/journal.ppat.1002510.

Martin, M. (2011). Cutadapt removes adapter sequences from high-throughput sequencing reads. EMBnet.Journal 17, 10–12. https://doi.org/10.14806/EJ.17.1.200.

Marzi, M.J., Ghini, F., Cerruti, B., De Pretis, S., Bonetti, P., Giacomelli, C., Gorski, M.M., Kress, T., Pelizzola, M., Muller, H., et al. (2016). Degradation dynamics of micrornas revealed by a novel pulse-chase approach. Genome Res. 26, 554–565. https://doi.org/10.1101/gr.198788.115.

Massirer, K.B., Perez, S.G., Mondol, V., and Pasquinelli, A.E. (2012). The miR-35-41 family of microRNAs regulates RNAi sensitivity in Caenorhabditis elegans. PLoS Genet. 8, e1002536. https://doi.org/10.1371/journal.pgen.1002536.

la Mata, M., Gaidatzis, D., Vitanescu, M., Stadler, M.B., Wentzel, C., Scheiffele, P., Filipowicz, W., and Großhans, H. (2015). Potent degradation of neuronal mi RNA s induced by highly complementary targets. EMBO Rep. 16, 500–511. https://doi.org/10.15252/embr.201540078.

McJunkin, K., and Ambros, V. (2014). The embryonic mir-35 family of microRNAs promotes multiple aspects of fecundity in Caenorhabditis elegans. G3 (Bethesda). 4, 1747–1754. https://doi.org/10.1534/g3.114.011973.

McJunkin, K., and Ambros, V. (2017). A microRNA family exerts maternal control on sex determination in C. elegans. Genes Dev. 31, 422–437. https://doi.org/10.1101/gad.290155.116.

Miki, T.S., Rüegger, S., Gaidatzis, D., Stadler, M.B., and Großhans, H. (2014). Engineering of a conditional allele reveals multiple roles of XRN2 in Caenorhabditis elegans development and substrate specificity in microRNA turnover. Nucleic Acids Res. 42, 4056–4067. https://doi.org/10.1093/nar/gkt1418.

Paix, A., Wang, Y., Smith, H.E., Lee, C.Y.S., Calidas, D., Lu, T., Smith, J., Schmidt, H., Krause, M.W., and Seydoux, G. (2014). Scalable and versatile genome editing using linear DNAs with microhomology to Cas9 sites in Caenorhabditis elegans. Genetics 198, 1347–1356. https://doi.org/10.1534/genetics.114.170423.

Parchem, R.J., Moore, N., Fish, J.L., Parchem, J.G., Braga, T.T., Shenoy, A., Oldham, M.C., Rubenstein, J.L.R., Schneider, R.A., and Blelloch, R. (2015). miR-302 Is Required for Timing of Neural Differentiation, Neural Tube Closure, and Embryonic Viability. Cell Rep. 12, 760–773. https://doi.org/10.1016/j.celrep.2015.06.074.

Piwecka, M., Glažar, P., Hernandez-Miranda, L.R., Memczak, S., Wolf, S.A., Rybak-Wolf, A., Filipchyk, A., Klironomos, F., Cerda Jara, C.A., Fenske, P., et al. (2017). Loss of a mammalian circular RNA locus causes miRNA deregulation and affects brain function. Science 357, eaam8526. https://doi.org/10.1126/science.aam8526.

Reichholf, B., Herzog, V.A., Fasching, N., Manzenreither, R.A., Sowemimo, I., and Ameres, S.L. (2019). Time-Resolved Small RNA Sequencing Unravels the Molecular Principles of MicroRNA Homeostasis. Mol. Cell 75, 756–768.e7. https://doi.org/10.1016/j.molcel.2019.06.018.

Rissland, O.S., Hong, S.-J., and Bartel, D.P. (2011). MicroRNA destabilization enables dynamic regulation of the miR-16 family in response to cell-cycle changes. Mol. Cell 43, 993–1004. https://doi.org/10.1016/j.molcel.2011.08.021.

Sherrard, R., Luehr, S., Holzkamp, H., McJunkin, K., Memar, N., and Conradt, B. (2017). Mirnas cooperate in apoptosis regulation during c. Elegans development. Genes Dev. 31, 209–222. https://doi.org/10.1101/gad.288555.116.

Sheu-Gruttadauria, J., Pawlica, P., Klum, S.M., Wang, S., Yario, T.A., Schirle Oakdale, N.T., Steitz, J.A., and MacRae, I.J. (2019). Structural Basis for Target-Directed MicroRNA Degradation. Mol. Cell 75, 1243–1255.e7. https://doi.org/10.1016/j.molcel.2019.06.019.

Shi, C.Y., Kingston, E.R., Kleaveland, B., Lin, D.H., Stubna, M.W., and Bartel, D.P. (2020). The ZSWIM8 ubiquitin ligase mediates target-directed microRNA degradation. Science 370, eabc9359. https://doi.org/10.1126/science.abc9359.

Shukla, S., Bjerke, G.A., Muhlrad, D., Yi, R., and Parker, R. (2019). The RNase PARN Controls the Levels of Specific miRNAs that Contribute to p53 Regulation. Mol. Cell 73, 1204–1216.e4. https://doi.org/10.1016/j.molcel.2019.01.010.

Stoeckius, M., Maaskola, J., Colombo, T., Rahn, H.-P., Friedländer, M.R., Li, N., Chen, W., Piano, F., and Rajewsky, N. (2009). Large-scale sorting of C. elegans embryos reveals the dynamics of small RNA expression. Nat. Methods 6, 745–751. https://doi.org/10.1038/nmeth.1370.

Tran, A.T., Chapman, E.M., Flamand, M.N., Yu, B., Krempel, S.J., Duchaine, T.F., Eroglu, M., and Derry, W.B. (2019). MiR-35 buffers apoptosis thresholds in the C. elegans germline by antagonizing both MAPK and core apoptosis pathways. Cell Death Differ. 26, 2637–2651. https://doi.org/10.1038/S41418-019-0325-6.

Vieux, K.-F., Prothro, K.P., Kelley, L.H., Palmer, C., Maine, E.M., Veksler-Lublinsky, I., and McJunkin, K. (2021). Screening by deep sequencing reveals mediators of microRNA tailing in C. elegans. Nucleic Acids Res. 49, 11167–11180. https://doi.org/10.1093/nar/gkab840.

Wang, Z., Hou, Y., Guo, X., vanderVoet, M., Boxem, M., Dixon, J.E., Chisholm, A.D., and Jin, Y. (2013). The EBAX-type Cullin-RING E3 Ligase and Hsp90 Guard the Protein Quality of the SAX-3/Robo Receptor in Developing Neurons. Neuron 79, 903–916. https://doi.org/10.1016/J.NEURON.2013.06.035.

Wu, E., Thivierge, C., Flamand, M., Mathonnet, G., Vashisht, A.A., Wohlschlegel, J., Fabian, M.R., Sonenberg, N., and Duchaine, T.F. (2010). Pervasive and cooperative deadenylation of 3’UTRs by embryonic microRNA families. Mol. Cell 40, 558–570. https://doi.org/10.1016/j.molcel.2010.11.003.

Wyman, S.K., Knouf, E.C., Parkin, R.K., Fritz, B.R., Lin, D.W., Dennis, L.M., Krouse, M.A., Webster, P.J., and Tewari, M. (2011). Post-transcriptional generation of miRNA variants by multiple nucleotidyl transferases contributes to miRNA transcriptome complexity. Genome Res. 21, 1450–1461. https://doi.org/10.1101/gr.118059.110.

Yang, A., Shao, T.-J., Bofill-De Ros, X., Lian, C., Villanueva, P., Dai, L., and Gu, S. (2020a). AGO-bound mature miRNAs are oligouridylated by TUTs and subsequently degraded by DIS3L2. Nat. Commun. 11, 2765. https://doi.org/10.1038/s41467-020-16533-w.

Yang, B., Schwartz, M., and McJunkin, K. (2020b). In vivo CRISPR screening for phenotypic targets of the mir-35-42 family in C. elegans. Genes Dev. 34, 1227–1238. https://doi.org/10.1101/gad.339333.120.

Ye Duan, Isana Veksler-Lublinsky, V.A. (2021). Critical contribution of 3’ non-seed base pairing to the in vivo function of the evolutionarily conserved let-7a microRNA. Biorxiv https://doi.org/10.1101/2021.03.29.437276.

Zanin, E., Dumont, J., Gassmann, R., Cheeseman, I., Maddox, P., Bahmanyar, S., Carvalho, A., Niessen, S., Yates, J.R., Oegema, K., et al. (2011). Affinity purification of protein complexes in C. elegans. Methods Cell Biol. 106, 289–322. https://doi.org/10.1016/B978-0-12-544172-8.00011-6.

Zeng, Y., Yi, R., and Cullen, B.R. (2005). Recognition and cleavage of primary microRNA precursors by the nuclear processing enzyme Drosha. EMBO J. 24, 138–148. https://doi.org/10.1038/sj.emboj.7600491.

Zhao, Y., Jin, L., Wang, Y., Kong, Y., and Wang, D. (2019). Prolonged exposure to multi-walled carbon nanotubes dysregulates intestinal mir-35 and its direct target MAB-3 in nematode Caenorhabditis elegans. Sci. Rep. 9. https://doi.org/10.1038/S41598-019-48646-8.

