## Supplemental Figure S1-S7 for "The developmentally-timed decay of an essential microRNA family is seed sequence-dependent"

**List of Supplemental Materials**

**Figure S1** miRNA Taqman qPCR standard curves used for absolute quantification of *mir-35* variants and *mir-36*

**Figure** **S2** An alternative design of *mir-35(seed_rev)* also attenuates embryo to L1 decay

**Figure S3** Summary of miRNA expression changes in *mir-35(seed_rev)* and *mir-35(seed_mut)*

**Figure S4** Two-phase regulation model and EBAX-1-mediated regulation of *mir-35-42*

**Figure S5** Summary of miRNA expression changes in *mir-35(mut_3')* and *mir-35(mir-82_3')*

**Figure S6** miRNA tailing and trimming of *mir-35* variants in L1

**Figure S7** Decay of *mir-35(mut_3')* isoforms at the embryo to L1 transition

**Table S1** Strains used in this study

**Table S2** Oligonucleotides used in this study and allele information

**Table S3** Samples used for deep sequencing

**Table S4** miRNA abundance and percent single-nucleotide tails in deep sequencing samples

**Table S5** Spike-in normalized reads from deep sequencing of staged wild type and *ebax-1(null)* embryos and L1 larvae

**Table S6** Analysis of oligonucleotide tails on *mir-35-42* and *mir-35* variants.

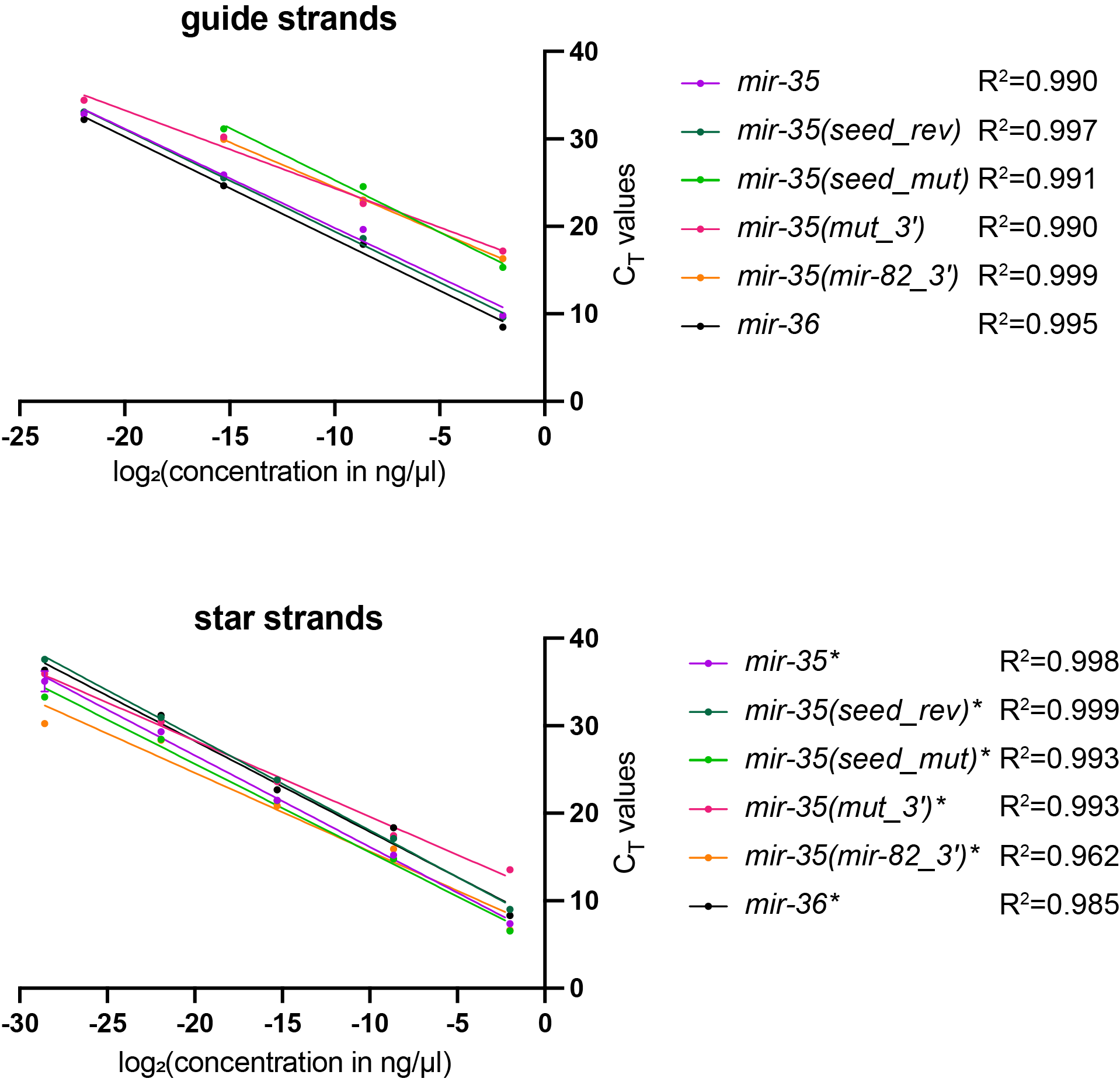

**Figure S1. miRNA Taqman qPCR standard curves used for absolute quantification of *mir-35* variants and *mir-36*** Mean and SEM for three technical replicates are shown (but error bars are smaller than the symbol in most cases). For guide strands, C_T_ values greater than 35 were excluded since this increased R^2^, and all experimental values still fell within truncated ranges. For star strands, all values were retained to allow for interpolation of all experimental values. Linear regression lines and R^2^ are shown.

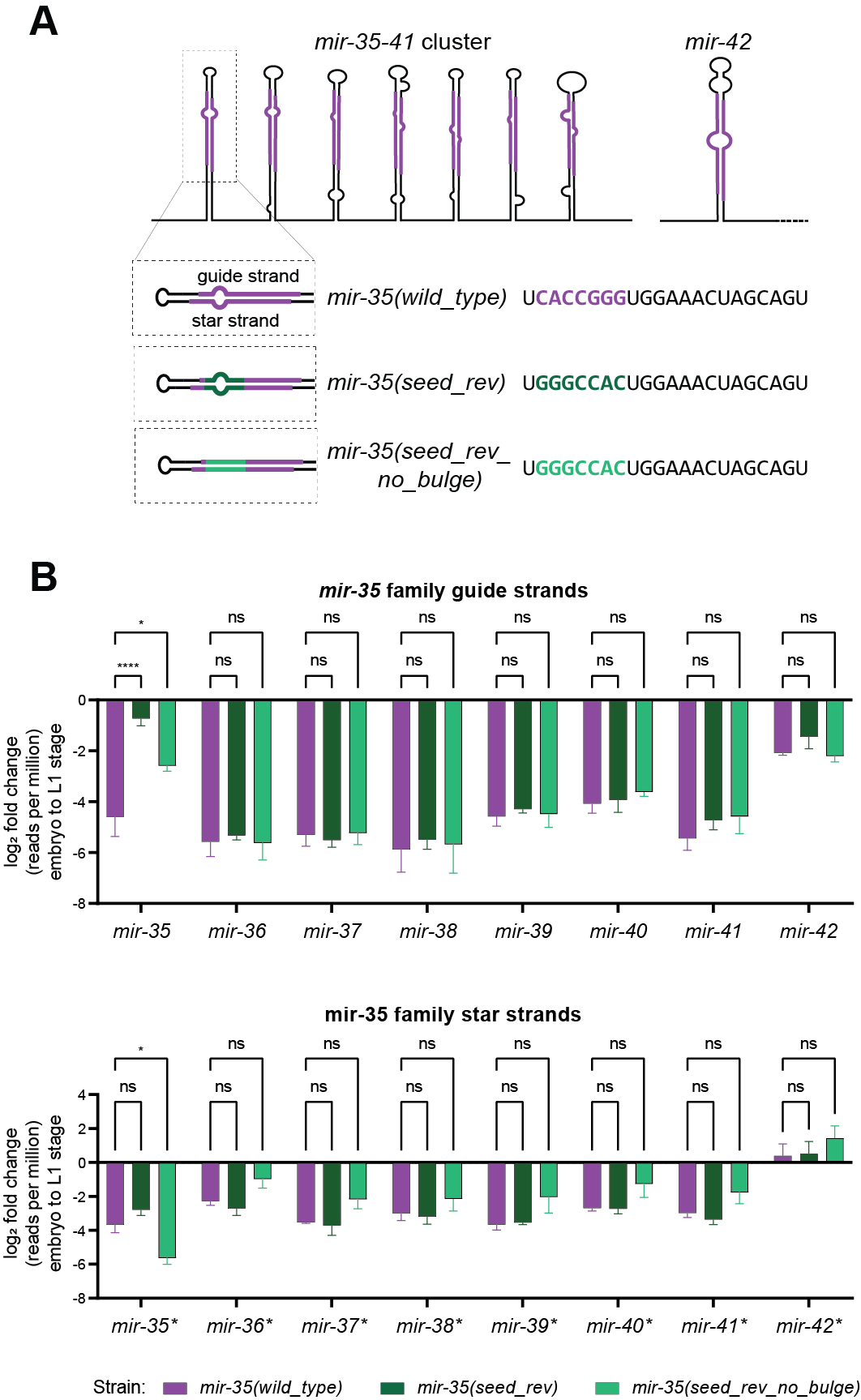

**Figure** **S2.** **An alternative design of *mir-35(seed_rev)* also attenuates embryo to L1 decay** A) A second allele generating a mature *mir-35* variant with a reversed seed sequence was generated. In this variant, bulges were removed from the precursor secondary structure. B) Log_2_(fold change) from embryo to L1, calculated from normalized reads per million in deep sequencing data. Like *mir-35(seed_rev)*, the decay of *mir-35(seed_rev_no_bulge)* from embryo to L1 was attenuated compared to wild type *mir-35.*

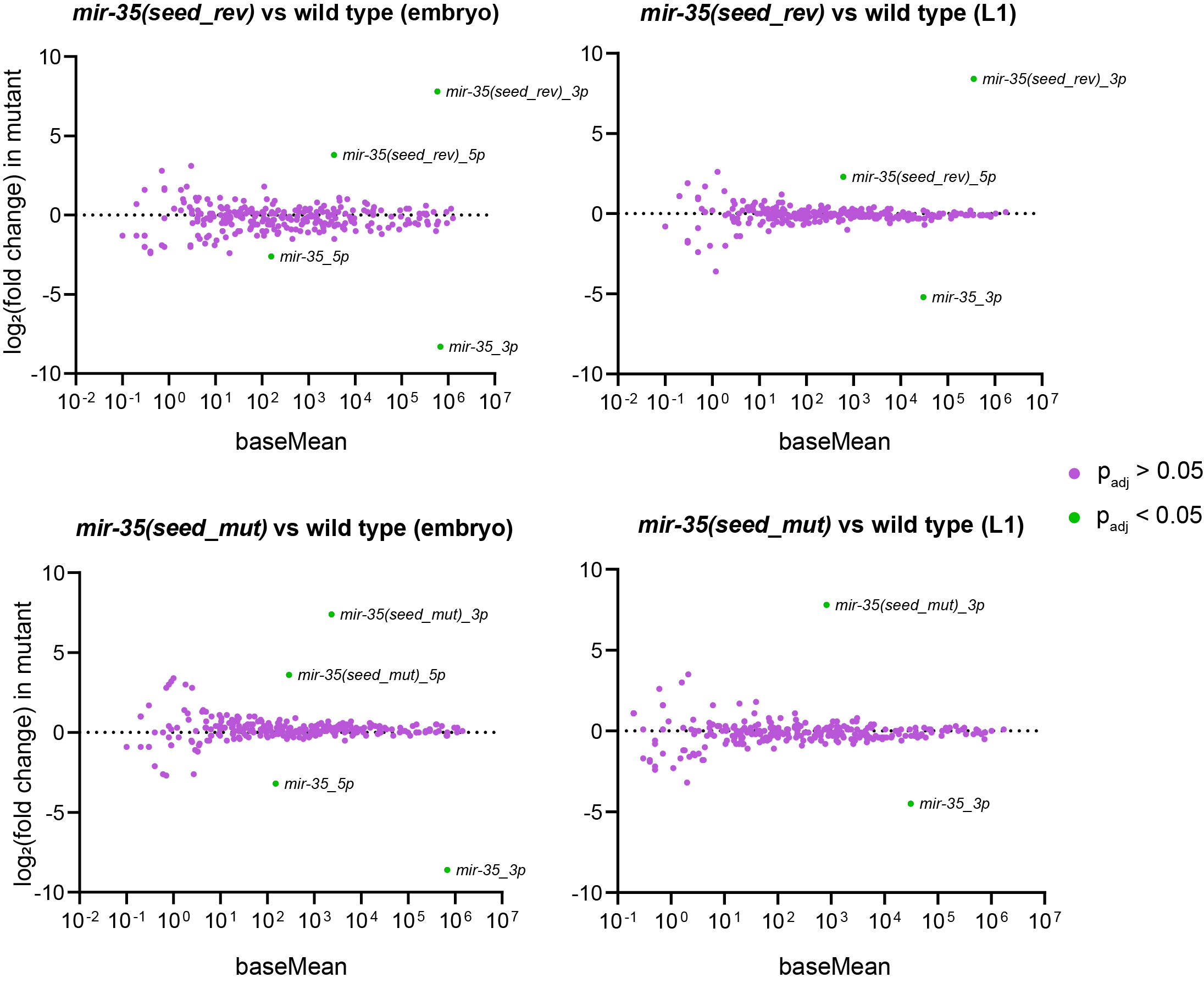

**Figure S3. Summary of miRNA expression changes in *mir-35(seed_rev)* and *mir-35(seed_mut)*** MA plots of deep sequencing results showing average abundance across all samples (baseMean) on x-axis and log_2_(fold change) in mutant versus wild type on y-axis. The only significant changes (green) are loss of wild type *mir-35* and gain of the mutant *mir-35* variant.

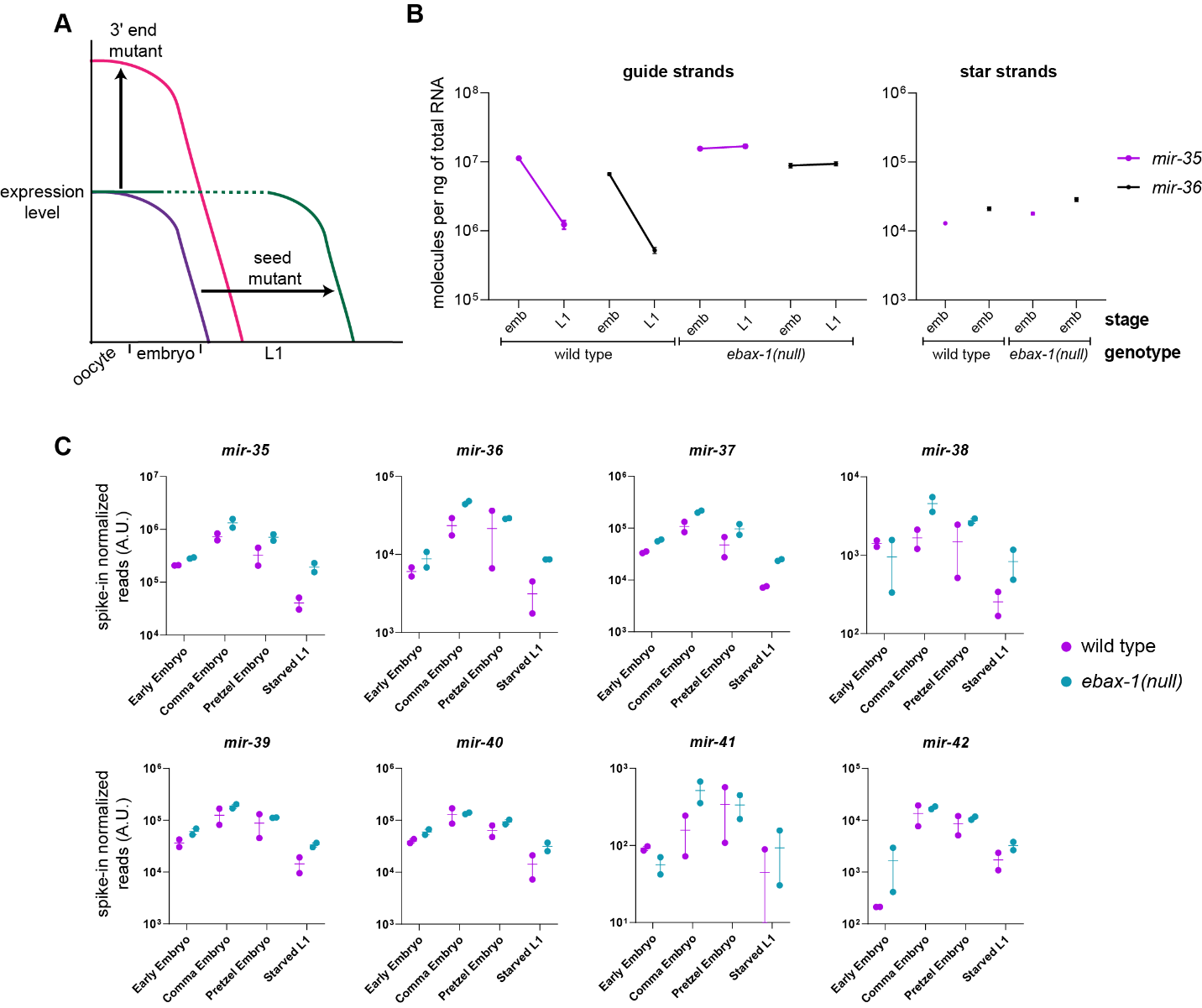

**Figure S4. Two-phase regulation model and EBAX-1-mediated regulation of *mir-35-42*** A) Mutations of the *mir-35* seed sequence strongly stabilize *mir-35* at the embryo to L1 transition, without increasing abundance in the embryo. Mutations to the 3' region result in elevated *mir-35* levels in the embryo, without disrupting decay at the embryo to L1 transition. This suggests that two distinct mechanisms may regulate *mir-35* abundance in these two developmental windows. B) Taqman-qPCR of *mir-35* and *mir-36* in wild type or *ebax-1(null)* mixed-stage embryo and L1 samples. Mean and SEM of four biological replicates shown. C) Deep sequencing of *mir-35-42* in wild type and *ebax-1(null)* hand-picked staged embryos and L1s. Spike-in RNA oligos were added prior to RNA isolation (see methods) and used to normalize miRNA reads. Two biological replicates shown with mean and SEM.

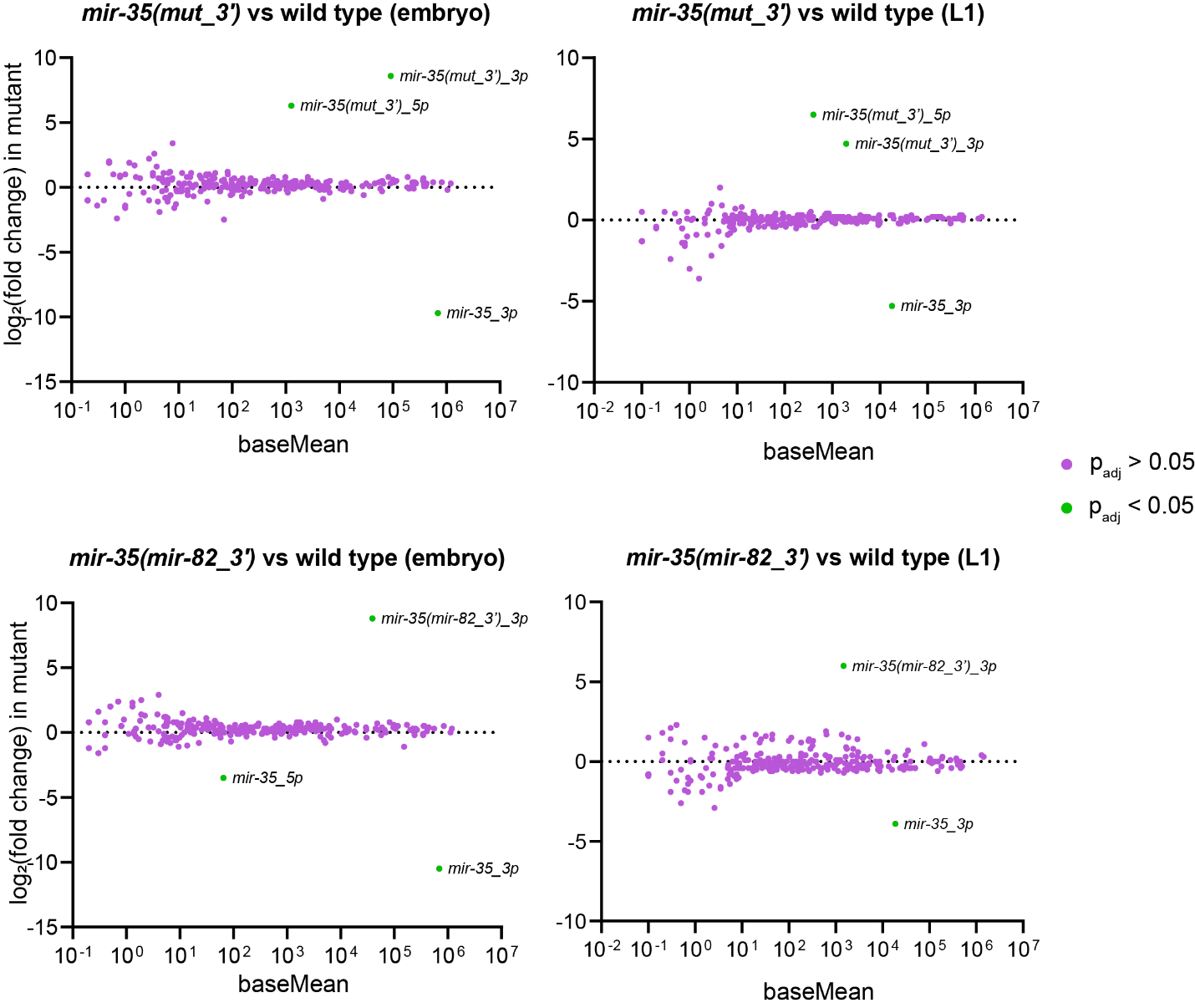

**Figure S5. Summary of miRNA expression changes in *mir-35(mut_3')* and *mir-35(mir-82_3')*** MA plots of deep sequencing results showing average abundance across all samples (baseMean) on x-axis and log_2_(fold change) in mutant versus wild type on y-axis. The only significant changes (green) are loss of wild type *mir-35* and gain of the mutant *mir-35* variant.

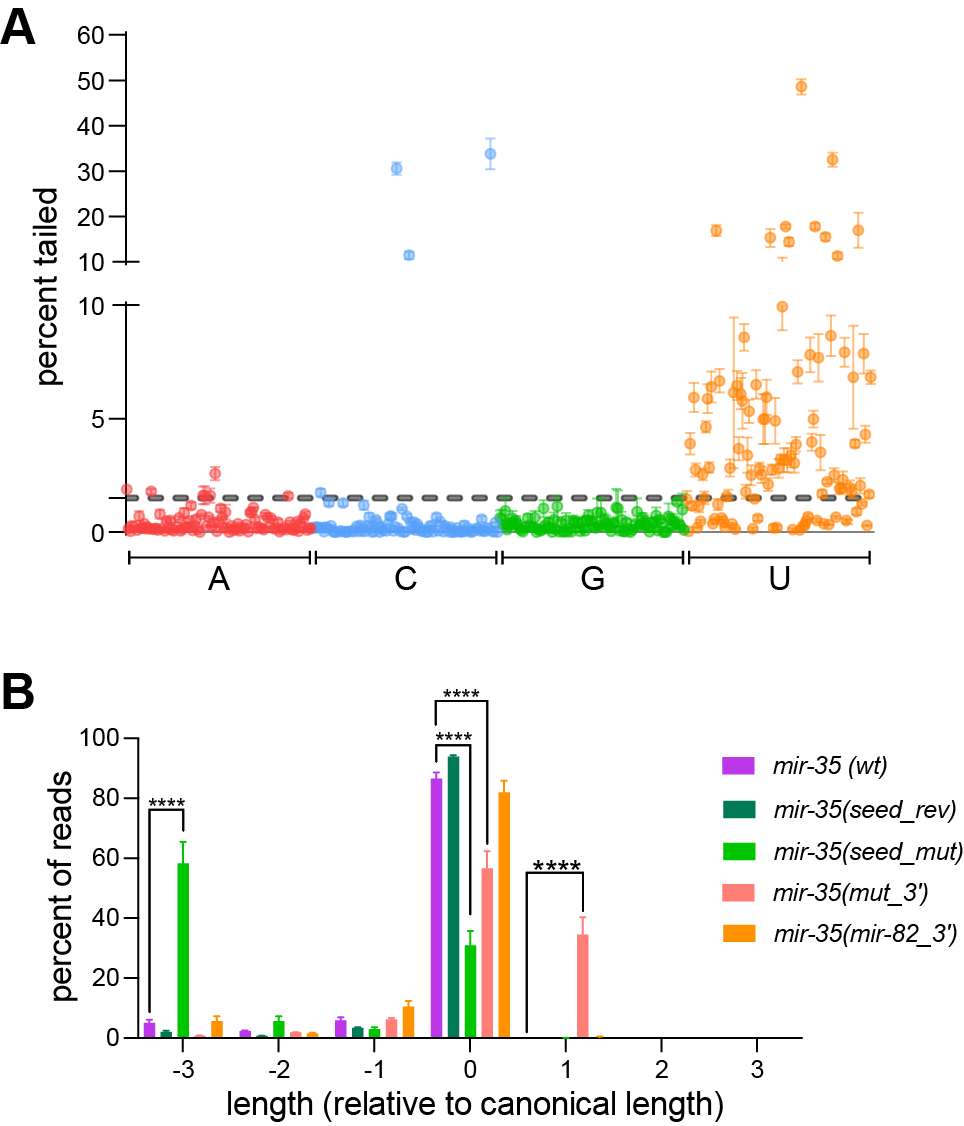

**Figure S6.** **miRNA tailing and trimming of *mir-35* variants in L1** A) Percent of reads with single nucleotide addition to the 3' end is shown for each miRNA with >50 RPM in L1. Mean and SEM are shown for six wild type replicates. (B) Percent of reads of each length (excluding tail), relative to the canonical length of *mir-35* in L1. Mean and SEM are shown for six replicates for wild type and three replicates for all *mir-35* variant strains. For each nucleotide, one-way ANOVA was performed, followed by Sidak’s multiple comparison test. ****p-value < 0.0001.

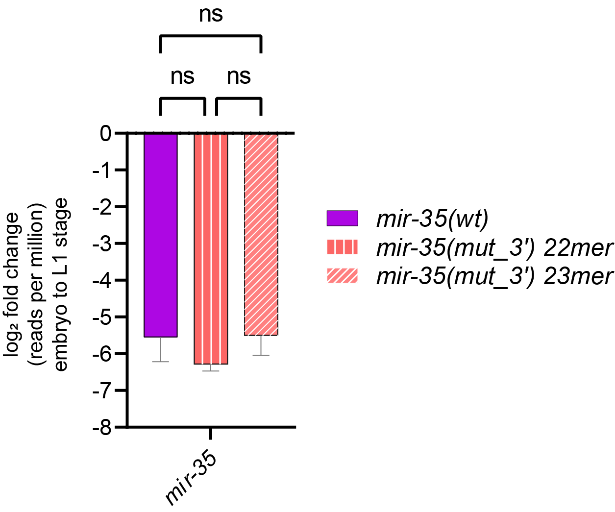

**Figure S7. Decay of *mir-35(mut_3')* isoforms at the embryo to L1 transition** Two prominent isoforms of *mir-35(mut_3')* guide strand are generated from the *mir-35(mut_3')* primary transcript. The decay of these two isoforms at the embryo to L1 transition is similar to each other and to wild type *mir-35.* Log_2_(fold change) was calculated from normalized reads from deep sequencing.
